# A Hollow TFG Condensate Spatially Compartmentalizes the Early Secretory Pathway

**DOI:** 10.1101/2024.03.26.586876

**Authors:** William R. Wegeng, Savannah M. Bogus, Miguel Ruiz, Sindy R. Chavez, Khalid S. M. Noori, Ingrid R. Niesman, Andreas M. Ernst

**Author notes:** These authors contributed equally to this work (William R. Wegeng, Savannah M. Bogus).

## Abstract

In the early secretory pathway, endoplasmic reticulum (ER) and Golgi membranes form a nearly spherical interface. In this ribosome-excluding zone, bidirectional transport of cargo coincides with a spatial segregation of anterograde and retrograde carriers by an unknown mechanism. We show that at physiological conditions, Trk-fused gene (TFG) self-organizes to form a hollow, anisotropic condensate that matches the dimensions of the ER-Golgi interface. Regularly spaced hydrophobic residues in TFG control the condensation mechanism and result in a porous condensate surface. We find that TFG condensates act as a molecular sieve, enabling molecules corresponding to the size of anterograde coats (COPII) to access the condensate interior while restricting retrograde coats (COPI). We propose that a hollow TFG condensate structures the ER-Golgi interface to create a diffusion-limited space for bidirectional transport. We further propose that TFG condensates optimize membrane flux by insulating secretory carriers in their lumen from retrograde carriers outside TFG cages.

## Main Text

In the early secretory pathway, the interface formed between the endoplasmic reticulum (ER) and the Golgi stack collaborates in the secretion of biosynthetic cargo^1^. Specialized domains of the ER, termed ER exit sites (ERES), form metastable contacts with the *cis* face of the Golgi apparatus. Morphologically, the ER-Golgi interface is defined by concave membranes at ERES and the cis-Golgi, resulting in a nearly spherical, ribosome-excluding zone < 500 nm in diameter^2,3^. The ER-Golgi interface functions as the primary ‘valve’ of the secretory pathway: in a typical human cell, ∼30% of the proteome as well as membrane lipids are routed through this interface, corresponding to ∼50% of the ER volume emptying into this junction every 40 minutes, while simultaneously retrieving ∼90% of the membrane and recycling it back to the ER^1,4^. The exchange of material at the interface is accomplished by bidirectional transport via COPII and COPI complexes, which deform the membrane to produce vesicular and tubular carriers. COPII carriers originate in the ER and are directed towards the Golgi (anterograde), while COPI carriers operate in the opposite (retrograde) direction^5^. The small, submicron dimension of the interface thus poses the problem of clashes between carriers headed in the opposite direction, which would result in content mixing and eventually loss of compartmental identity. Several decades ago, immuno-EM approaches revealed a strict spatial separation of COPI and COPII machineries, with COPII carriers concentrated in the center of the interface while COPI coats were segregated towards the periphery^6–9^. How this spatial separation is achieved remains unexplained.

Recently, evidence has emerged suggesting condensation of the ER-Golgi interface proteins TANGO1 at ERES, Sec16 and Trk-fused gene (TFG) within the interface, and GM130 at the cis-Golgi^10–18^, suggesting that the early secretory pathway is structured by self-organizing protein collectives. Among these proteins, Sec16 and TFG have been proposed as candidates capable of structuring the space between ERES and the cis-Golgi. Sec16, a peripheral membrane protein, was found to co-condense with TANGO1^18^, which is a transmembrane component of ERES^12^. Importantly, the heterogeneous Sec16 condensates were found to impact cargo flux through the early secretory pathway. Silencing of TFG in *C. elegans* strongly impacted ER-Golgi interface morphology and resulted in both diminished Golgi membranes and mislocalized COPII carriers^10^, which is paralleled by results obtained from HeLa cells that support a role for TFG in organizing ERES and anterograde carriers into larger, coherent structures^19^. Similar to Sec16, TFG was shown to critically impact the amount of cargo exported from the ER^10^, but not its export kinetics, while secretion of bulky cargo (i.e. collagen) was significantly impaired^19^. When recombinant TFG purified from bacteria was subjected to high concentrations of potassium acetate (KOAc), it precipitated out of solution to form amorphous, reversible aggregates^11^. These data prompted the speculation that TFG might form a biomolecular condensate capable of organizing the ER-Golgi interface as ‘molecular glue’, with additional functions in uncoating of anterograde carriers^13,14^. How a putative TFG condensate could contribute to the spatial segregation of anterograde and retrograde carriers at the interface remained elusive.

### Recombinant TFG assembles into anisotropic, hollow condensates in vitro

We set out to test whether a protein condensate at the ER-Golgi interface simultaneously defined its structure and controlled the organization of bidirectional traffic. If the intrinsically disordered proteins Sec16 and TFG were responsible for structuring the spherical ER-Golgi interface, they should localize between ERES and cis-Golgi membranes. To localize TFG and Sec16 at individual ERES/Golgi pairs, we employed the microtubule-destabilizing agent nocodazole to ‘unlink’ the convoluted Golgi ‘ribbon’^20^. Next, we co-transfected cells with mGFP-Sec16L and FLAG-TFG-SNAP, and immunolabeled cis-Golgi (GM130) and ERES (TANGO1) markers. Localization analysis indicated that TFG was present precisely between ERES and the cis-Golgi, whereas Sec16 appeared restricted to the ERES membranes (**Fig. 1a; Extended Data Fig. 1a, b**).

**Figure 1.**
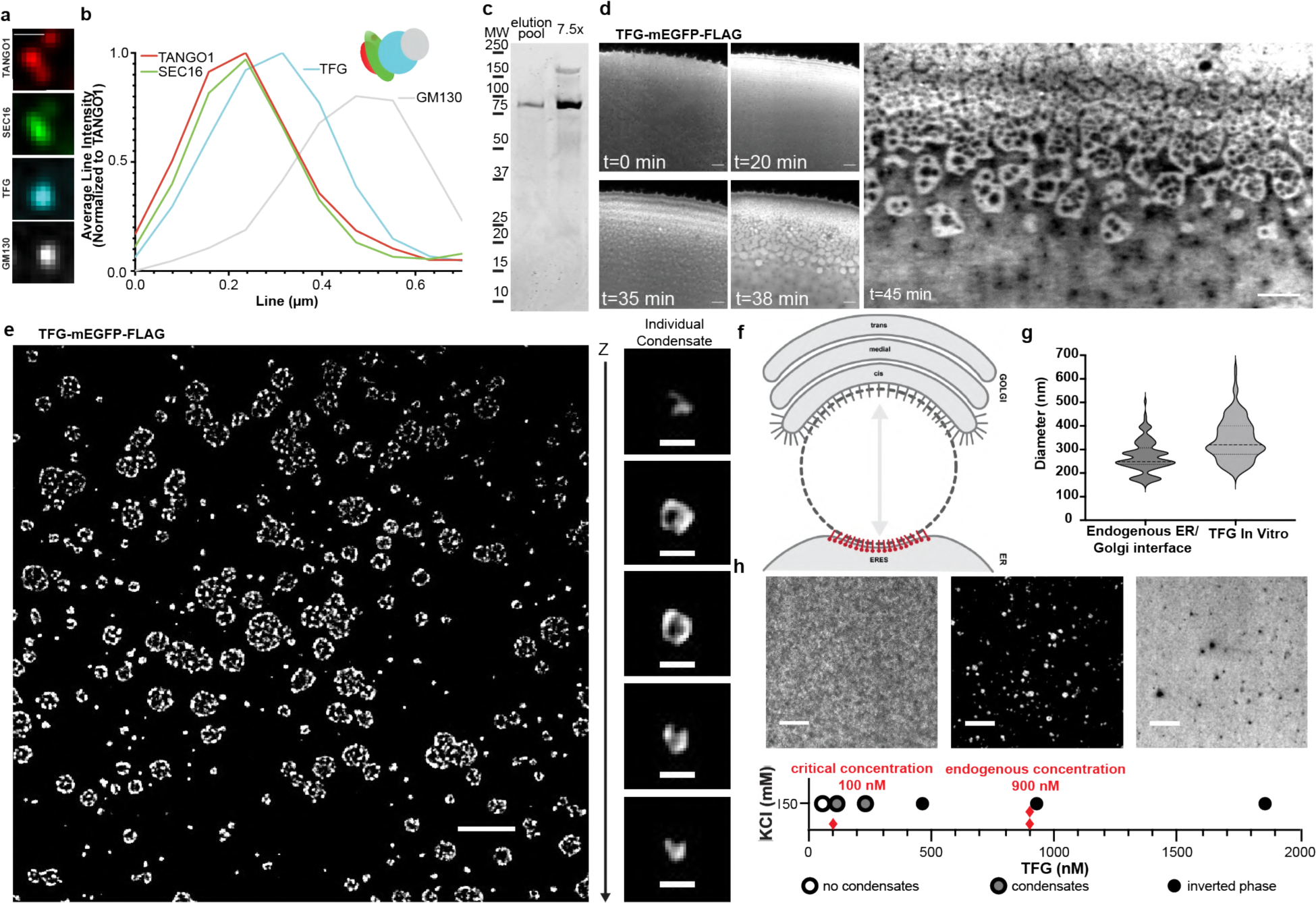
Recombinant TFG assembles into anisotropic, hollow condensates *in vitro*. **a** Representative 4-color micrograph of an individual ERES/cis-Golgi unit. HeLa cells were transfected with mGFP-Sec16L, FLAG-TFG-SNAP (labeled with SNAP-Cell 647-SiR), and immunostained for endogenous GM130 and TANGO1. **b** Fluorescence intensity line scan of 4-color ERES/cis-Golgi unit (averaged and normalized to TANGO1, n = 4). Schematic of localizations detected in representative unit (a) is given. **c** Representative purity of peak TFG elutions from concentration/salt exchange (Coomassie staining). **d** Confocal time series of increasing concentration of TFG-mEGFP-FLAG at the edge of an evaporating 10 µl sessile droplet, 150 mM KCl. Scale bar 5 μm. **e** TFG-mEGFP-FLAG spontaneously assembles into ‘sponge-like’, anisotropic condensates (HEPES/KOH pH 7.3; 150 mM KCl; 20% (v/v) PEG 8 kDa; confocal microscopy). Scale bar 5 µm. Individual TFG condensate, Z-step 125 nm. Scale bar 500 nm. **f** Schematic depicting the ER-Golgi interface (targets for immunolabeling at ERES: green; cis-Golgi: magenta. **g** ERES to cis-Golgi distance measured between immunolabeled GM130 and TANGO1, mean = 269, n = 120, vs. diameter of individual hollow TFG condensates *in vitro,* mean = 341, n = 175. **h** Determination of the critical concentration required for the formation of hollow TFG condensates (in the absence of crowding agents). Representative micrographs depicting the distribution of TFG below and above the critical concentration of 100 nM. Higher concentrations of TFG yield inverted phases (right micrograph). Red diamonds indicate critical concentration and average cellular concentration of TFG. Scale bars 5 µm.

Due to TFG populating the interface, we next set out to test if recombinant TFG was capable of forming condensates *in vitro*. For this purpose, we generated pure (>95%) recombinant TFG from human suspension cells (**Fig. 1c; Extended Data Fig. 1c**). When characterizing recombinant TFG in supersaturated solutions^15,21^, we concentrated the protein by sessile droplet evaporation and observed phase-separation into condensates with an anisotropic structure, forming ∼500 nm voids within the dense phase that paralleled the structure of condensates formed upon overexpression in cells (**Fig. 1d**). The anisotropic distribution of protein with these condensates developed from an initially isotropic distribution of TFG (**Fig. 1d**). These data support the sufficiency of TFG to form anisotropic condensates independently of other components.

We next tested whether formation of TFG condensates is promoted or inhibited by a crowding agent which is frequently employed *in vitro* to mimic macromolecular crowding experienced by proteins in the cytosol^22^. When recombinant TFG was exposed to PEG (8 kDa, 20% (v/v)), we observed rapid induction of TFG condensates with a broad size distribution (**Fig. 1e**). TFG formed both micron-sized, sponge-like condensates as well as condensates < 500 nm that appeared spherical, closely matching the dimensions of the ER-Golgi interface obtained from immunolabeling ERES and cis-Golgi markers (endogenous ER-Golgi interface: 270 nm ± 65; TFG condensates *in vitro*: 340 nm ± 87; **Fig. 1g**). The submicron condensates exhibited a void in their center, resembling the shape of a hollow sphere (**Fig. 1e**). Notably, the formation of anisotropic, hollow individual condensates did not result from the presence of nucleic acids in the condensate core (**Extended Data Fig. 1d**). These data suggested that the sponge-like appearance of larger condensates resulted from coalescence of individual <500 nm hollow condensates. Thus, TFG self-organizes to form submicron-sized, hollow spherical condensates that could explain the spherical void at the ER-Golgi interface. The micron-sized ‘sponge-like’ condensates may also represent physiologically relevant structures, as ERES are clustered under the Golgi ribbon^2^, a process that would be facilitated by the coalescence of individual submicron condensates. We next set out to test whether the submicron condensates would form under physiological conditions and concentrations (**Fig. 1h**). We concentrated recombinant TFG of defined starting concentrations stepwise to determine the critical concentration required for condensate formation. Condensation was observed already at concentrations of ∼100 nM, which is below the critical concentration typically reported for intrinsically disordered proteins to undergo liquid-liquid phase separation (LLPS; 10-100 µM^23–26^). Condensation at such a low critical concentration is consistent with specific interactions among TFG molecules. When approaching concentrations of ∼1 µM, we observed the formation of inverted phases with submicron aqueous inclusions. As the average cellular concentration of TFG is ∼900 nM^27^, these observations support the ability of TFG to form condensates under physiological conditions.

To confirm the physiological relevance of TFG as a cytosolic extension of ERES and its role in controlling flux of cargo from the ER, we employed the Retention Using Selective Hooks (RUSH^28^) system with a KDEL-Str hook and the model cargo SBP-eGFP-ECadherin to compare the rate of cargo export in wild type cells and cells in which expression of TFG was silenced. The absence of TFG significantly diminished the rate of cargo clearance from the ER and resulted in compromised cell growth (**Extended Data Fig. 2d**). This parallelled prior data obtained for the temperature-sensitive model cargo ts045-VSV-G, collagen, and TFG hepatocyte knockout mice^10,19,29^.

### Endogenous tagging and overexpression confirms assembly of TFG into anisotropic condensates that disassemble during mitosis

Next, we set out to test whether the hollow TFG condensates observed *in vitro* could be detected in cells at physiological concentrations. Employing CRISPR/Cas9, we endogenously tagged TFG at its C-terminus with mClover-FLAG, and validated heterozygous clones by sequencing and Western blot (**Extended Data Fig. 3b**). When performing live-cell microscopy of TFG::mClover-FLAG, we observed discrete foci reminiscent of condensates throughout the cell, as well as a diffuse distribution in the cytosol supportive of the presence of a two-phase coexistence(i.e., dilute and dense TFG phases) (**Fig. 2a**; **Extended Data Fig. 3e**). In the dense TFG foci, the distribution appeared to be anisotropic, paralleling our results obtained *in vitro* (**Fig. 1e**). Furthermore, the average diameter of TFG condensates *in vivo* closely mirrored the dimensions observed *in vitro* (TFG condensates *in vitro*: 341 nm ± 87; TFG condensates *in vivo*: 293 nm ± 87), strongly supporting that TFG can self-organize into hollow condensates that match the dimensions of the ER-Golgi interface in live cells and at physiological concentrations and conditions. When photobleaching entire endogenous TFG condensates *in vivo*, we observed a low extent of fluorescence recovery (FRAP). Analysis of the mobility of TFG in endogenous condensates (partial FRAP) could not be robustly performed due to their submicron dimension and dynamics.

**Figure 2.**
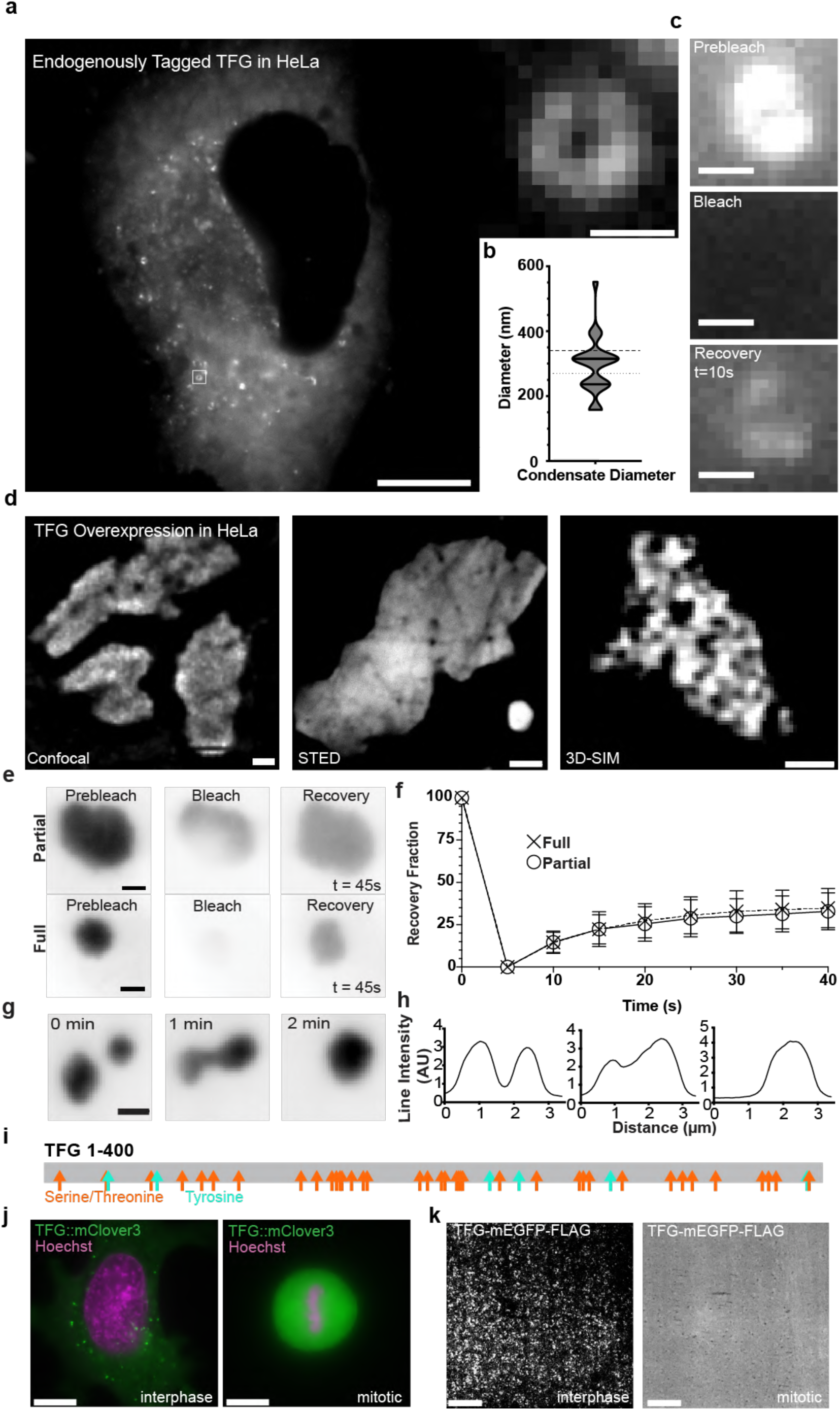
Endogenous tagging and overexpression confirms assembly of TFG into anisotropic condensates that disassemble during mitosis. **a** Representative widefield micrograph of HeLa cells with endogenously tagged TFG::mClover-FLAG. Scale bar 10 μm. Magnified micrograph (dashed box in cell overview): individual TFG condensate. Scale bar 500 nm. **b** Quantification of the diameter of endogenous TFG condensates. n = 25. Dashed line represents average of *in vitro* condensate diameter and dotted line represents average of endogenous ER/Golgi interface diameter, for reference. **c** Qualitative fluorescence recovery after photobleaching (FRAP) of a TFG condensate. Scale bar 500 nm. **d** Left: Live-cell confocal microscopy of HeLa cells transfected with TFG-mEGFP-FLAG. Scale bar 5 μm. Middle: STED micrograph of HeLa cells transfected with TFG-mEGFP-FLAG. Scale bar 2 μm. Right: 3D-Structured Illumination Microscopy (3D-SIM) of HeLa cells transfected with TFG-tGFP. Scale bar 500 nm. **e** Representative micrographs of partial and full FRAP experiments on overexpressed TFG condensates in HeLa cells. **f** Quantification of average condensate recovery rates for partial (n = 33) and full (n = 30) FRAP; normalized to pre-bleach (= 100) and post-bleach intensity (= 0). **g** Representative live-cell imaging of TFG condensate fusion dynamics. **h** Line scan of condensate depicted in g. **i** Schematic depicting putative phosphorylation sites within TFG: serine/threonine, orange; tyrosine, cyan. Putative phosphorylation sites were predicted by NetPhos-3.0. **j** Representative micrographs of interphase (left) and mitotic (right) HeLa cells with endogenously tagged TFG. Cells were stained with Hoechst (magenta). Scale bar 10 μm. k *In vitro* analysis of recombinant TFG-162-240-mEGFP-FLAG purified from synchronized mitotic or interphase Expi293F cells. Scale bar 10 μm.

To further characterize the dynamics of TFG condensates, we overexpressed TFG in HeLa cells. TFG initially exhibited a dispersed localization throughout the cytosol, and suddenly began to condense into dynamic sub-micron foci (**Extended Data Fig. 3a**). These foci increased in size over the scale of minutes and fused to form structures several microns large. Condensation of TFG upon overexpression did not depend on the mEGFP tag, since it was observed with diverse genetically encoded fluorescent (SNAP, mCherry) and epitope tags (**Extended Data Fig. 3c**). Furthermore, TFG condensates efficiently recruited mGFP-Sec16L to their surface (**Extended Data Fig. 3d**), paralleling reports of a specific interaction between Sec16 and TFG^10^. To gain further insight into the distribution of TFG within condensates, we employed both stimulated emission depletion (STED) and 3D-structured illumination (3D-SIM) super-resolution microscopy (**Fig. 2d**). TFG-mEGFP-FLAG condensates neither exhibited smooth surfaces nor a high degree of sphericity (**Fig. 2d**; **Extended Data Fig. 4b**), but rather appeared sponge-like, paralleling our results obtained from supersaturated TFG and upon crowding *in vitro*. Interestingly, the voids within the dense condensate phase exhibited diameters that resemble the diameter of the ER-Golgi interface, mirroring the *in vitro* observation that sponge-like condensates result from coalescence of individual hollow TFG condensates (**Fig. 1e**). FRAP experiments supported a low mobility of TFG within condensates, as well as a low extent of exchange of TFG between the dense condensate phase and cytosol (partial FRAP: n=33; full FRAP: n=30; **Fig. 2e, f**) indicating viscoelastic material properties, with approximately 70% of TFG molecules’ diffusion being significantly confined within the condensate dense phase. However, micron-sized TFG condensates exhibited ‘capillary bridges’ during fusion and relaxed back into overall spherical geometries after fusion, which together indicate the presence of surface tension and the formation of flexible TFG collectives (**Fig. 2g, h**). When performing live-cell confocal imaging of large TFG condensates, we observed diffraction-limited heterogeneities supportive of an anisotropic distribution of TFG within the condensate dense phase (**Extended Data Fig. 4b**). Our data thus far highlight TFG’s propensity to self-organize into ER-Golgi interface-sized, anisotropic (hollow) condensates. However, the low mobility of TFG within the dense phase raised the question how these structures could be regulated and disassembled. Multiple proteins that populate the early secretory pathway are regulated by phosphorylation^18,30,31^, leading to the dispersal of ERES and Golgi membranes upon mitotic entry, and followed by a rapid reassembly upon mitotic exit^32^. TFG contains several putative serine/threonine as well as tyrosine phosphorylation sites that are dispersed across its entire length (**Fig. 2i**), suggesting that condensates could be regulated by phosphorylation as well. We thus set out to test whether TFG would disassemble during mitosis *in vivo* and *in vitro*. Capitalizing on the endogenously tagged TFG::mClover-FLAG cell line, we identified mitotic cells via Hoechst staining (**Fig. 2j**). Strikingly, TFG condensates could not be detected in mitotic cells, instead exhibiting a dispersed localization within the cytosol. We next set out to validate this result *in vitro* by synchronizing Expi293F cells prior to transfection and overexpression of TFG-mEGFP-FLAG. Mirroring the results obtained from endogenously tagged TFG, recombinant TFG purified from mitotic cells failed to form condensates, while protein purified from interphase cells efficiently assembled into hollow condensates (**Fig. 2k**). These data support that TFG is dynamically regulated across the cell cycle along with other structural proteins in the early secretory pathway. We propose that phosphorylation sites across the protein may lead to repulsion and thus inhibition of condensate formation, with dephosphorylation allowing rapid reassembly of TFG into hollow spheres.

### Hydrophobic ‘sticker’ residues control the anisotropic condensation mechanism of TFG

We next set out to identify which regions within TFG were responsible for the formation of hollow condensates (**Fig. 3a**). TFG was reported to assemble into homooctamers via an amino-terminal PB1 domain^11^, and is predicted by AlphaFold^33^ to have two additional amphipathic alpha helices (**Extended Data Fig. 5a, b**). However, the majority of TFG is intrinsically disordered (73%; **Extended Data Fig. 5c**) and harbors a Q-rich region in which every third amino acid is glutamine. We purified recombinant TFG fragments (**Extended Data Fig. 6a-f**) that represent the folded moiety (TFG*1-123*), the initial segment of IDR (IDR1: TFG*124-240*; fragments within IDR1: *124-161* and *162-240*), the Q-rich tract (IDR2: TFG*241-359*), as well as the short carboxy-terminal fragment (IDR3: TFG*360-400*), and tested for the ability of each fragment to form anisotropic condensates (**Fig. 3b**). While the folded moiety of TFG (TFG*1-123*) failed to form condensates, all fragments within the intrinsically disordered moiety of TFG formed anisotropic, hollow condensates. Although the average diameter of fragment condensates largely matched the diameter observed for full-length TFG condensates, significant differences were detected in the size distribution among fragments (**Fig. 3c**; full-length TFG*1-400* condensates: 341 nm ± 87 SD, n = 175; diameter range of TFG fragments: 290-400 nm). We next performed FRAP experiments on condensates formed by TFG fragments and compared them to condensates formed by full-length TFG (**Extended Data Fig. 6g, h**). While condensates formed by the carboxy-terminal moiety of TFG (Q-rich TFG*241-359* and TFG*360-400*) approached mobile fractions of ∼25%, both full-length and individual amino-terminal fragments of TFG exhibited reduced mobility, which matched the low mobility observed within condensates formed by full-length TFG *in vivo*. TFG interacts with Sec16 via its amino-terminus^34^ and TFG condensates are coated by Sec16 in cells (**Extended Data Fig. 3d**), suggesting that the amino-terminus of TFG faces the cytosol while its carboxy-terminus points to the condensate lumen. Furthermore, such an orientation would be consistent with a role proposed for the carboxy-terminus of TFG in COPII carrier uncoating^13^. These data provide evidence that multiple regions within the intrinsically disordered moiety of TFG can support the formation of anisotropic condensates.

**Figure 3.**
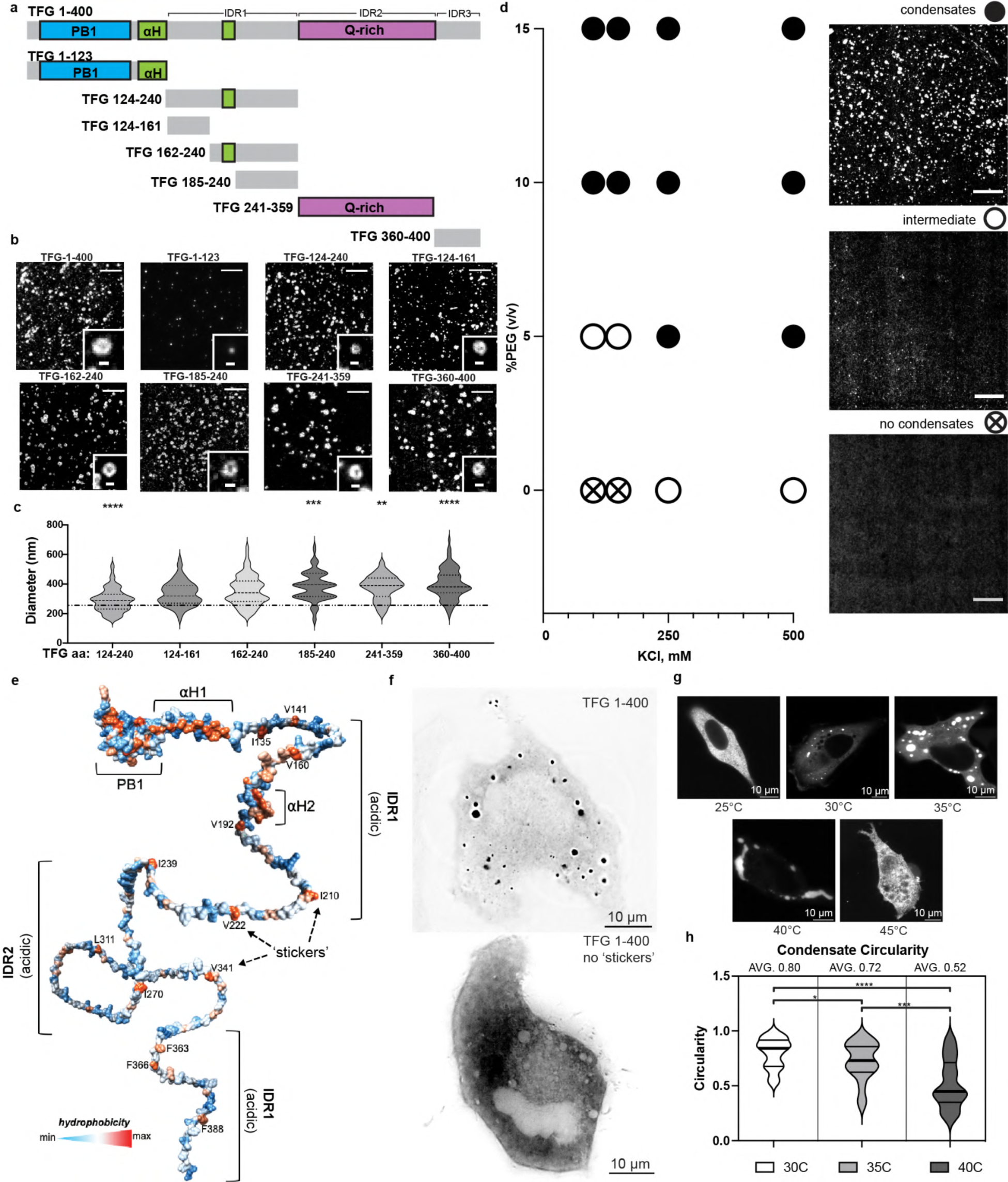
Hydrophobic ‘sticker’ residues control the anisotropic condensation mechanism of TFG. **a** Schematic depicting domains present within TFG: PB1 domain, blue; alpha helices, green; Gln-rich region, purple; intrinsically disordered region (IDR), grey. Individual IDRs are defined as: IDR1 (net acidic), IDR2 (Gln-rich), IDR 3 (net basic). Residues for truncated constructs are depicted to scale. **b** All truncated constructs except for residues 1-123 of TFG support the formation of lumen-containing (anisotropic) condensates. Representative micrographs are given. Overview scale bars 5 µm; insets 500 nm. **c** Quantification of the average peak-to-peak diameter of condensates formed by individual truncations of TFG. TFG; TFG 124-240: n = 124; TFG 124-162: n = 79; TFG 163-240: n = 176; TFG 241-359: n = 93; TFG 360-400: n = 107. Dashed line represents the average TFG 1-400 condensate diameter for reference. Two-tailed unpaired t-tests (****: p<0.001, ***: p<0.001, **: p<0.05). **d** TFG phase diagram probing increasing concentrations of KCl versus PEG 8 kDa as a crowding agent. TFG-mEGFP-FLAG: 70 nM; HEPES/KOH pH 7.3; X mM KCl as indicated; X% (v/v) PEG 8 kDa as indicated. Scale bar 10 µm. **e** Schematic based on an AlphaFold prediction of TFG structure color-coded for hydrophobicity (red: max; blue: min). Domain annotations and the position of hydrophobic residues (‘stickers’) within IDRs are given. **f** Widefield deconvolution micrographs of HeLa cells transfected with TFG-mEGFP-FLAG and TFG-1-400-w/o ‘stickers’-mEGFP-FLAG, respectively. Scale bar 10 µm. **g** Representative micrographs of HeLa cells transfected with TFG-mEGFP-FLAG shifted to the indicated temperatures. Scale bar 10 µm. **h** Quantification of condensate circularity at specific temperatures.

We next set out to test whether electrostatic or hydrophobic interactions contribute to the anisotropic condensation mechanism by screening full-length TFG against increasing volume fractions of crowding agent and salt (**Fig. 3d**; **Extended Data Fig. 7**). TFG more readily condensed in the presence of elevated salt concentrations, i.e., screening out electrostatic interactions, pointing to the involvement of hydrophobic residues in the formation of hollow condensates. Strikingly, within IDRs 1-3, TFG harbors hydrophobic residues that exceed the typical spacing observed in coiled-coil proteins, and which are present in all truncations capable of forming anisotropic condensates (**Fig. 3e**). This organization was highly reminiscent of the recently proposed ‘stickers and spacers’ model that describes the mechanistic basis for phase separation of intrinsically disordered proteins^35^ and suggested why individual fragments of TFG were able to support the formation of anisotropic condensates. The differential spacing of hydrophobic residues within TFG fragments may further cause the differences in size observed across condensates formed by TFG fragments. Importantly, cells transfected with full-length TFG in which hydrophobic ‘sticker’ residues were deleted, condensates failed to form entirely *in vivo*, further highlighting the importance of these residues in the condensation mechanism (**Fig. 3f**). The relative strengths of ‘sticker-sticker’ interactions were shown to depend on their spacing across IDRs, as well as temperature^35,36^. Thus, we set out to probe whether TFG condensates obtained from overexpression would depend on temperature by shifting the cells from 37°C to defined temperatures (**Fig. 3g**). In cells shifted below 30°C, TFG condensates disassembled and dispersed in the cytosol, while shifting the cells to 40°C promoted condensate coalescence. Strikingly, condensates at 30°C were nearly spherical, with an increase in temperature resulting in a successive reduction in condensate sphericity concomitant with an anisotropic, sponge-like appearance (**Fig. 3h**). These data suggest that a sticker-driven condensation mechanism controls assembly of TFG into hollow condensates, with interaction strengths among sticker residues maximizing near physiological temperatures.

Consistent with the ability of TFG to condense, we observed that overexpression of TFG diminished the amount of endogenous ERES and resulted in disassembly and fragmentation of the Golgi ribbon (**Extended Data Fig. 8a**). This suggests that endogenous TFG is sequestered into TFG condensates formed by overexpression, which would mimic the phenotype of TFG silencing and result in the previously reported defects in the organization of the early secretory pathway^10,19^. Strikingly, if ‘sticker’ residues are deleted in full-length TFG, endogenous ERES and Golgi markers remain unaffected, which strongly supports the specificity of TFG interactions via ‘sticker’ residues. Additionally, when both *wildtype* and ‘sticker’-lacking variants of TFG are co-overexpressed, TFG condensates diminish with increasing concentrations of ‘sticker’-lacking TFG (**Extended Data Fig. 8b**), which further supports that homotypic TFG interactions depend on hydrophobic ‘sticker’ residues.

### Hollow TFG condensates are porous and select for macromolecules of defined sizes

We next set out to further characterize TFG condensates by scanning electron microscopy (SEM) employing the anisotropic condensate-forming fragment TFG*162-240*. Condensates appeared to have a uniform surface, suggesting enclosure of an aqueous lumen by a network of TFG. We next performed transmission electron microscopy (TEM) to obtain high-resolution images of TFG condensates in the absence of fixatives or crowding agents (**Fig. 4a**). Strikingly, the TEM analysis revealed the presence of irregular, < 10 nm ‘pore-like’ openings on the surface of condensates, which prompted the hypothesis that proteins and cofactors sufficiently small could access the lumen of hollow TFG condensates. Next, we set out to test the relationship between molecular size and ability to access the lumen of TFG condensates by incubating TFG*162-240* condensates with fluorescent dextran species of increasing sizes (**Fig. 4c**, **Extended Data Fig. 9**). Dextrans smaller than 250 kDa readily accessed the condensate lumen, while larger dextrans were excluded – a feature that was preserved in condensates formed by full-length TFG (**Extended Data Fig. 9**) A cutoff of 250 kDa corresponds approximately to a protein diameter of < 8 nm^37^, which could explain why the ER-Golgi interface excludes ribosomes^3,38^ that are ∼20 nm in diameter.

**Figure 4.**
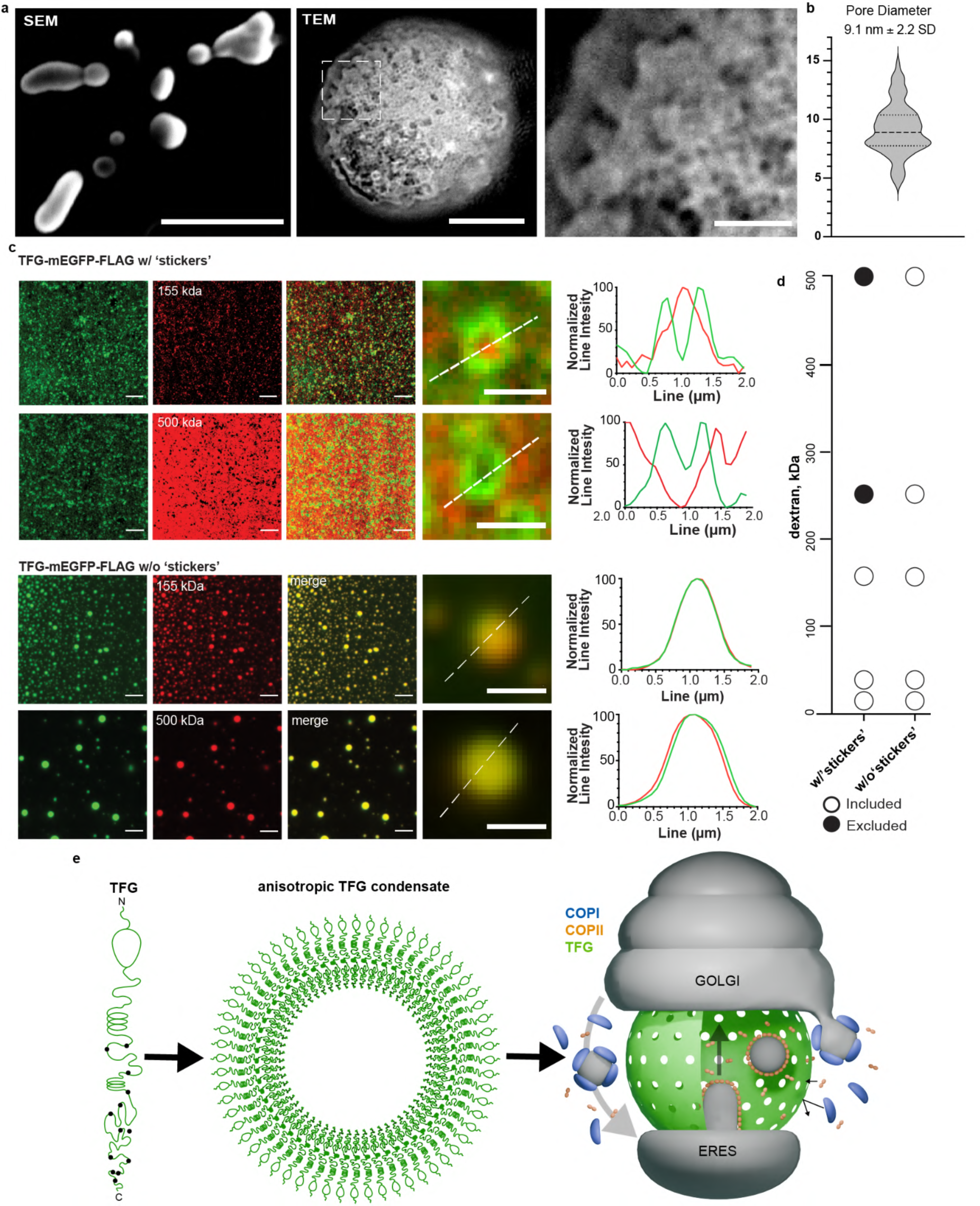
Hollow TFG condensates are porous and select for macromolecules of defined sizes. **a** Left: Scanning electron microscopy (SEM) of TFG-162-240-mEGFP-FLAG condensates. Scale bar 500 nm. Middle: transmission electron micrograph (TEM) of a single TFG-162-240-mEGFP-FLAG condensate (no crowding agents or negative staining; LUT inverted). Scale bar 100 nm. Right: Magnification of the porous surface of TFG condensates (boxed area in middle panel, inverted LUT). Scale bar 25 nm. **b** Quantification of average pore size of TFG condensates micrograph (n = 79). **c** Permeability of TFG-162-240-mEGFP-FLAG and TFG-185-240-w/o ‘stickers’ -mEGFP-FLAG condensates (induced with 20 % (v/v) PEG 8 kDa) for fluorescent dextran-TMR species of various sizes. Representative confocal micrographs are given for 155 kDa and 500 kDa dextran. Normalized line intensities for representative merged images are given. **d** Graph depicts molecular weight range of dextrans that can access the lumen of TFG condensates (included: detected in the lumen of TFG condensates, white-filled circles; excluded: sequestered from TFG condensates, back circles.) **e** Model depicting the role of TFG at the ER-Golgi interface: TFG self-organizes into 300-nm condensates that are hollow. Their porous surface allows for COPII coat components (Sec23/24, Sec13/31, both complexes <250 kDa) to access the lumen of TFG condensates, forming anterograde carriers. The retrograde carrier-forming COPI coat (550 kDa total) cannot access the lumen of TFG condensates. This results in a spatial segregation of anterograde from retrograde traffic, and the creation of a diffusion-limited space for bidirectional transport at the ER-Golgi interface.

We next purified a TFG fragment in which hydrophobic ‘stickers’ were removed (TFG185-240-w/o ‘stickers’). Upon crowding, highly dynamic, spherical condensates formed that lacked the anisotropic distribution of TFG. Furthermore, these condensates failed to exclude dextrans of higher molecular weights and lacked a porous surface when analyzed by TEM. (**Fig. 4c, Extended Data Fig. 10**). Thus, the porous structure and selective permeability of TFG condensates depends on its hydrophobic ‘sticker’ residues. Residues 174-184 encode for an amphipathic, alpha helical segment. While this region was not required for the formation of hollow condensates, did not affect the average size of openings on the condensate surface, nor impacted selectivity for dextrans (**Extended Data Fig. 10**), its presence significantly improved yields and reduced condensate hardening, pointing to a role in stabilizing lateral interactions among TFG molecules.

It was previously speculated that TFG could condense to form a ‘molecular glue’ that populates the ER-Golgi interface and mediates adhesion between ERES and the Golgi^14^, while promoting the uncoating of anterograde carriers^13^. We find that TFG indeed forms condensates at physiological conditions via hydrophobic ‘sticker’ residues, and assembles into hollow, porous spheres that act as molecular ‘sieves’. At the ER-Golgi interface, membrane carriers are routed in opposing directions at high rates^1^. While anterograde COPII carriers require the coat protein complexes Sec23/Sec24 and Sec13/Sec31 (each stable heterodimers < 250 kDa), retrograde COPI carriers require coatomer (a heptameric complex > 500 kDa) that is recruited *en bloc* to membranes^5,39,40^. Our data suggest that COPII coats could readily access the lumen of TFG condensates, while COPI coats would be excluded due to their size, leading to a spatial segregation of anterograde from retrograde transport at the ER-Golgi interface – i.e., anterograde carriers are contained within the lumen of hollow TFG condensates, while retrograde carriers are excluded from their lumen (**Fig. 4e**).

### Conclusion

Our findings address two long-standing questions regarding the anatomical and functional organization of the early secretory pathway. *First*, TFG self-organizes to form anisotropic, 300-nm hollow condensates that serve as cytosolic extensions of ERES, thereby structuring and controlling the dimensions of the ER-Golgi interface and putatively contributing to membrane concavity at both ERES and cis-Golgi membranes, both representing anatomical features that are conserved across multiple taxa^2,3,10^. This further results in TFG creating a diffusion-limited space between the ER and Golgi and may contribute to the overall polarization of the early secretory pathway. *Second*, based on discontinuities present on the surface of hollow TFG condensates, bidirectional transport at the ER-Golgi interface may be spatially segregated to ensure an efficient exchange of cargo between the organelles. This would prevent an aberrant diffusion of anterograde carriers away from target Golgi membranes, while avoiding collisions between uncoated anterograde and retrograde carriers. TFG condensates acting as molecular ‘sieves’ would be supported by the exclusion of ribosomes from the ER-Golgi interface and the segregation of vesicle-forming machinery (with COPII coats localizing to the interface, and COPI coats segregated to its periphery). Once assembled in the condensate interior, the COPII-coated carriers formed from Sec23/Sec24 and Sec13/31 heterodimers would exceed the exclusion size of the TFG boundary, leading to their concentration at the ER-Golgi interface. The resulting coalescence of anterograde carriers within the lumen of hollow TFG condensates could contribute to the formation of the ER-Golgi intermediate compartment (ERGIC)^42^, with energy-dependent docking and fusion machinery overcoming the TFG boundary to mediate content delivery to target membranes^43,44^. In agreement with these points, when TFG is depleted, the symmetry of the ER-Golgi interface is lost, resulting in an accumulation of vesicles outside the interface and a reduction of Golgi membranes concomitant with a reduced flux from the ER^10,19^. Notably, TFG condensates serving as extensions of ERES would be compatible with recently proposed ‘tunnels’ that may enable export of bulky cargo^17,41^. Importantly, our results are also compatible with recent reports that propose self-organization of transmembrane and peripheral ER-Golgi interface components^12,15,16,18^. Altogether, these findings lead to an emerging view in which the early secretory pathway is structured by protein collectives capable of dictating its morphology, connectivity, and performance.

## Acknowledgements

We thank Dr. Arshad Desai, Dr. Elizabeth Villa, Dr. Enfu Hui, Dr. Ishier Raote, Dr. Ünal Coskun, and Dr. Aleksander Rebane for discussions and feedback on the manuscript.

## Author contributions

W.R.W., S.M.B., M.R., S.R.C., and K.S.M.N. analyzed data and performed experiments. I.R.N. performed electron microscopy. A.M.E. conceived of the study and designed research. W.R.W., S.M.B., M.R., S.R.C., and A.M.E. wrote the manuscript.

## Data and materials availability

All source data and materials are available upon request.

## Extended Data Figures

**Extended Data Fig. 1.**
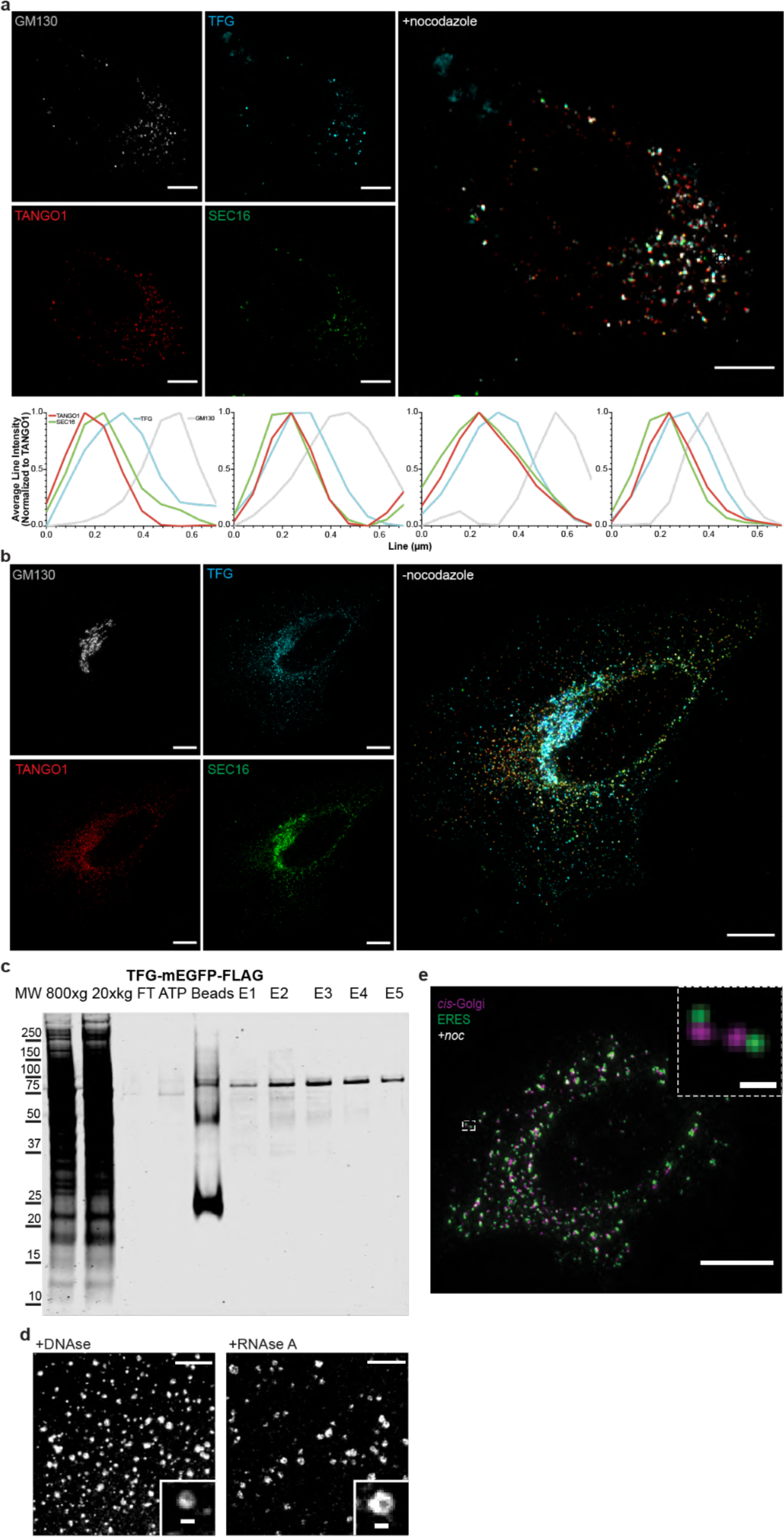
Localization of Sec16 and TFG at the ER-Golgi interface (cell overview for magnification provided in Fig. 1. **a** Micrographs of HeLa cells treated with nocodazole **a** (+noc) or **b** (-noc), transfected with mGFP-Sec16L (green), FLAG-TFG-SNAP (cyan) and immunolabeled for endogenous GM130 (gray) and TANGO1 (red), Scale bar 10 µm. Individual line scans of ER-Golgi units used for averaged line scan (Fig. 1b) are given. **c** Representative purification overview of TFG-mEGFP-FLAG from Expi293F suspension cells (Coomassie staining). Peak elution purity >95%. **d** TFG-162-240-mEGFP-FLAG lumen-containing condensates were treated with 1 U DNAse and imaged after a 3 minute incubation period. TFG-162-240-mEGFP-FLAG was purified in the presence of 10 U RNAse A (added during a wash step). (HEPES/KOH pH 7.3; 150 mM KCl; 20% (v/v) PEG 8 kDa). Scale bars 5 μm, 500 nm insets. **e** Micrograph of HeLa cells immunolabeled with ERES (green) and cis-Golgi (magenta) markers (max. intensity Z-projections). The Golgi ribbon was unlinked using nocodazole to facilitate visualization of individual ER-Golgi interfaces (+noc). Scale bar 10 µm, inset 500 nm.

**Extended Data Fig. 2.**
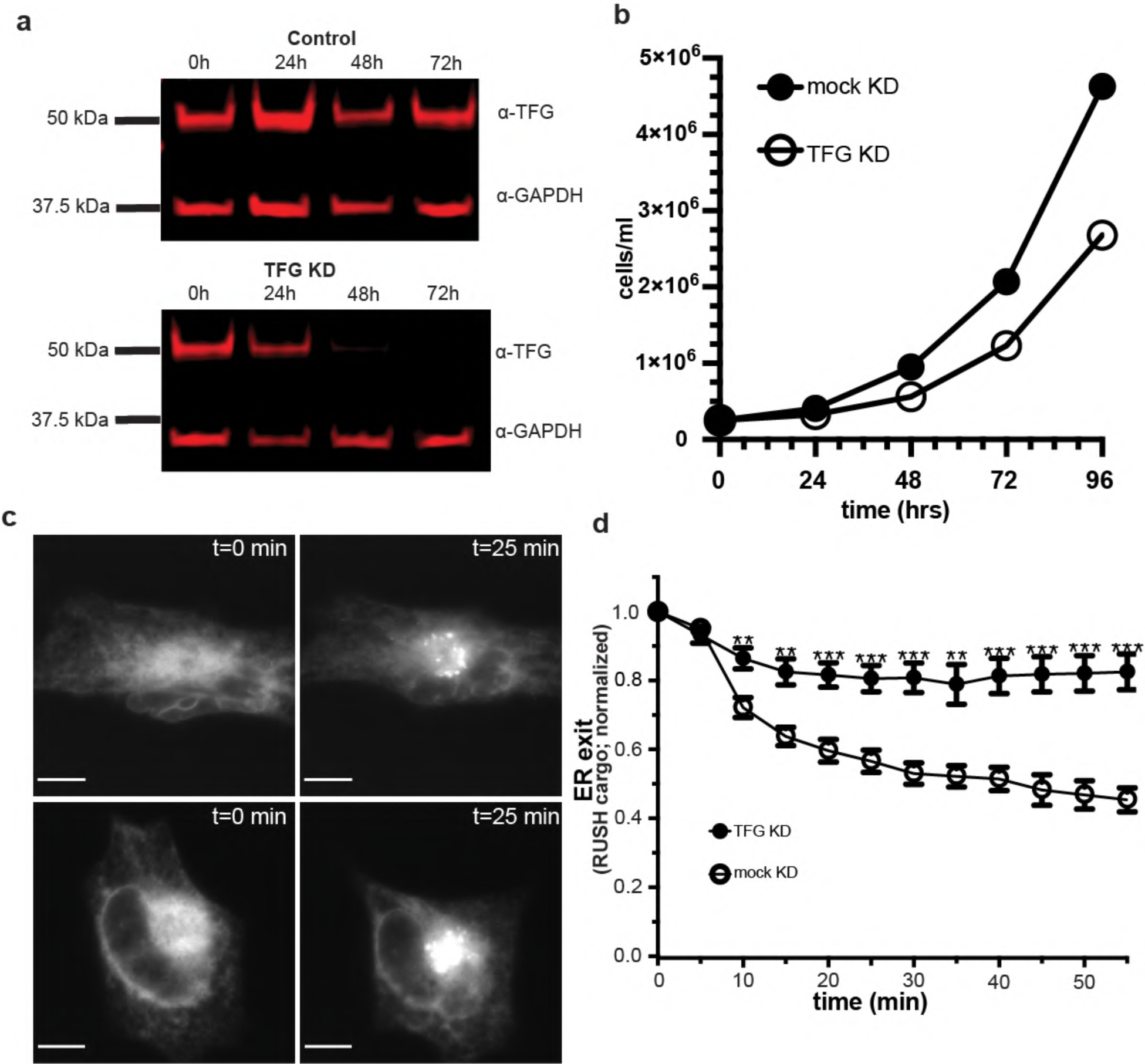
TFG impacts cargo export and cell growth. **a** Western blot of TFG siRNA knockdown and mock knockdown at 24, 48, and 72 hours. **b** Growth curve of TFG knockdown and mock knockdown Expi293F cells. **c** Live-cell visual correlates for RUSH assay. Cells were either subjected to mock or TFG siRNA treatment for 72 h and transfected with Str-KDEL_SBP-EGFP-ECadherin. Cargo waves triggered upon incubation with biotin; scale bar 10 µm. **d** Quantification of ER exit of RUSH cargo (ECadherin) in live cells for TFG knockdown (72 h; n = 9) and mock knockdown (72 h; n = 8) cells.

**Extended Data Fig. 3.**
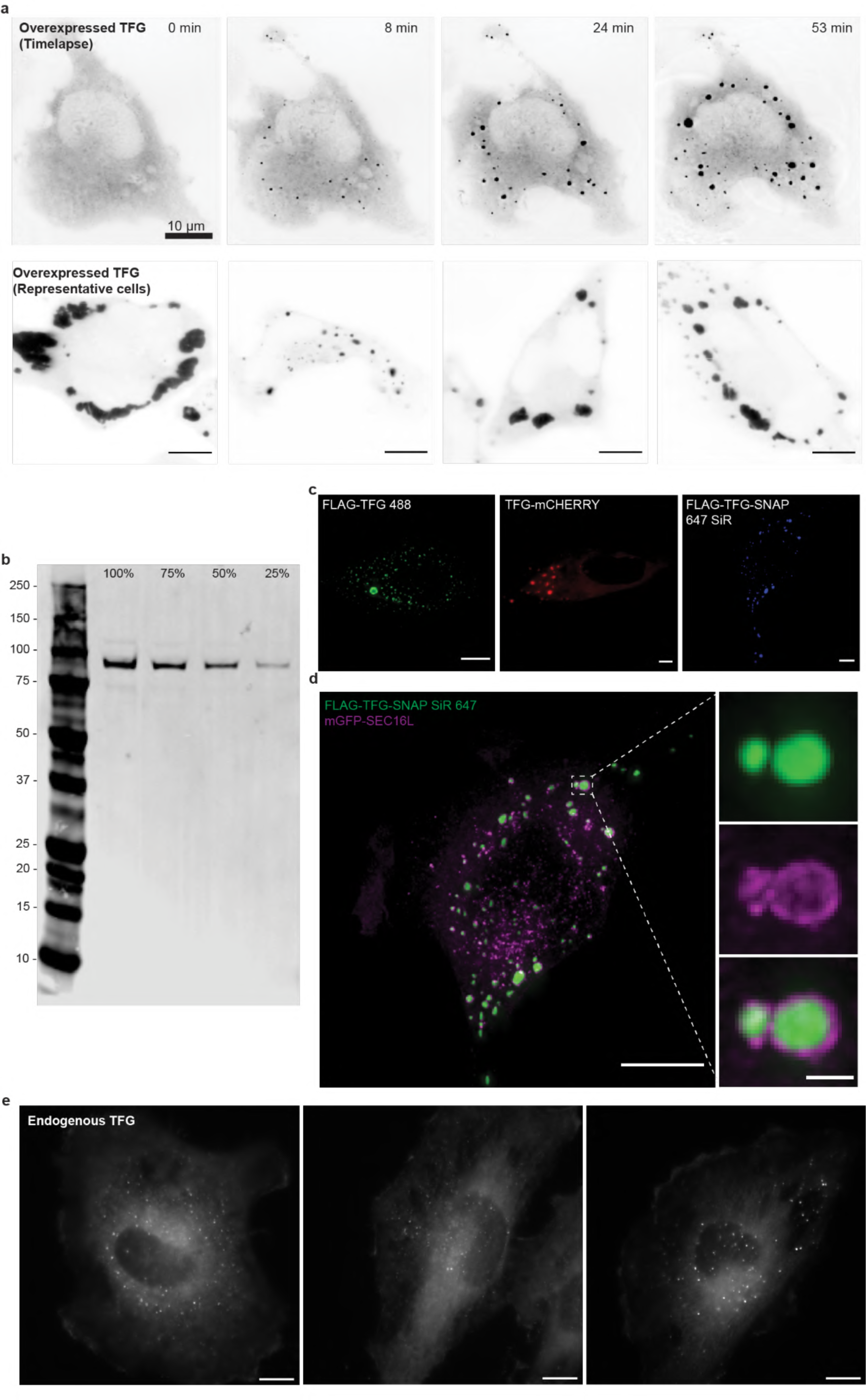
TFG condensates formed from overexpression are observed with a variety of tags and recruit Sec16 to their surface. **a** Top panel: Live-cell imaging of Hela cells transfected with TFG-mEGFP-FLAG 24 h post-transfection (widefield deconvolution microscopy, max. intensity Z-projections, intervals chosen to show nucleation and growth). Bottom panel: Representative images of TFG-mEGFP-FLAG overexpressed in HeLa cells exhibiting various condensate morphologies. **b** Anti-TFG Western blot of HeLa TFG::mClover-FLAG cells. 100%, 75%, 50%, 25% of samples loaded from lysates containing 70,000 cells. **c** Left panel: cells transfected with FLAG-TFG immunostained with anti-TFG, Alexa Fluor 488 and imaged with widefield deconvolution microscopy. Middle panel: live confocal image of cells transfected with TFG-mCherry-FLAG. Right panel: widefield deconvolution image of cells transfected with FLAG-TFG-SNAP and incubated with SNAP 647-SiR. Scale bar 5 µm. **d** Widefield deconvolution microscopy of HeLa cells co-transfected with mGFP-Sec16L and FLAG-TFG-SNAP. Left panel: Cell overview. Scale bar 10 μm. Right panel: magnification of TFG condensate; individual channels and merged image are given. Scale bar 5 μm. **e** Examples of HeLa cell clones harboring endogenously tagged TFG::mClover-FLAG. Scale bar 10 µm.

**Extended Data Fig. 4.**
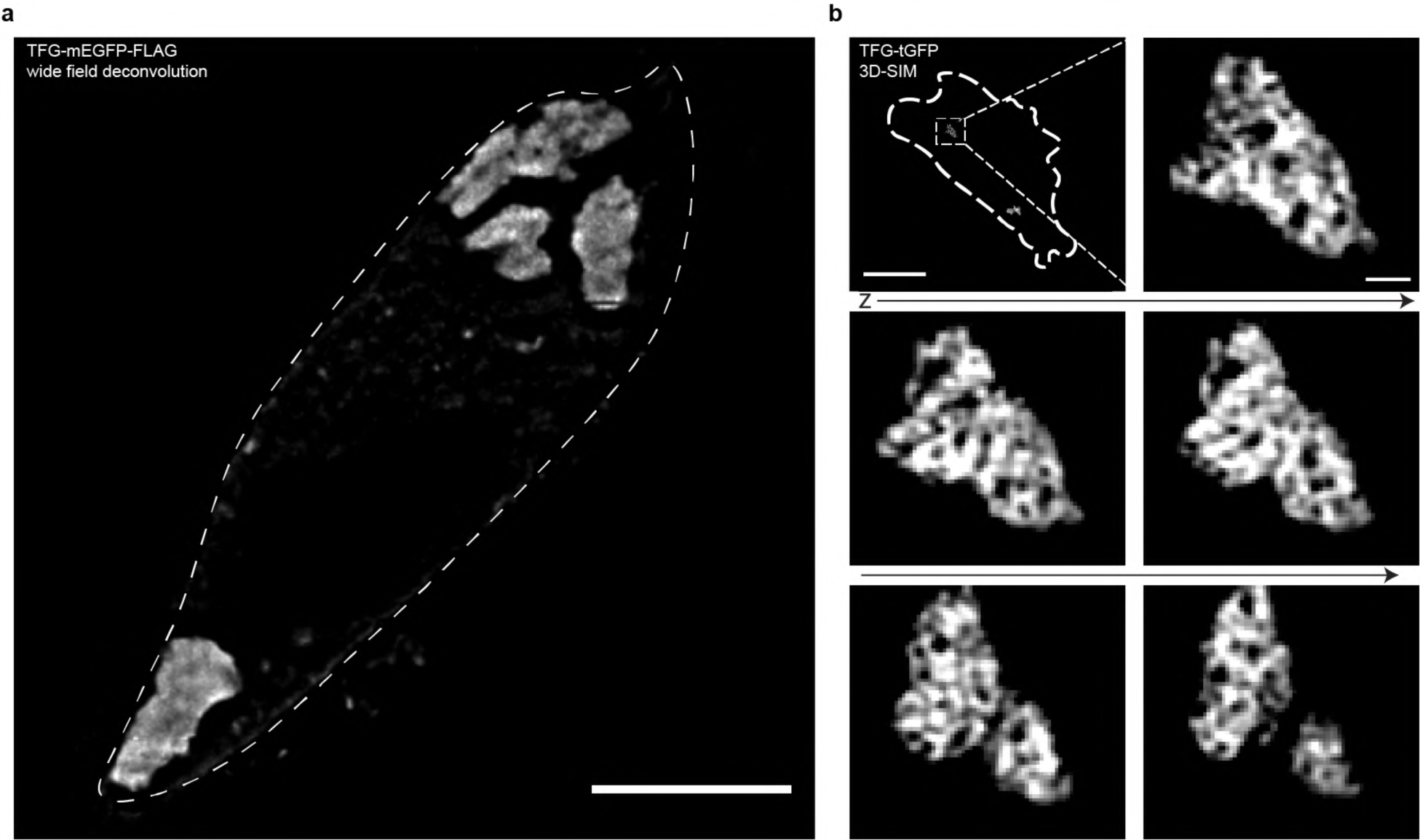
TFG condensates exhibit an anisotropic distribution in cells. **a** Live-cell widefield deconvolution microscopy of HeLa cells transfected with TFG-mEGFP-FLAG. Scale bar 10 μm. **b** 3D-Structured Illumination Microscopy (3D-SIM) of HeLa cells transfected with TFG-tGFP. Cell overview is provided (thick dashed line indicates cell outline), and serial sections are depicted. Z-step = 125 nm. Scale bar 10 µm. Magnification scale bar 500 nm.

**Extended Data Fig. 5.**
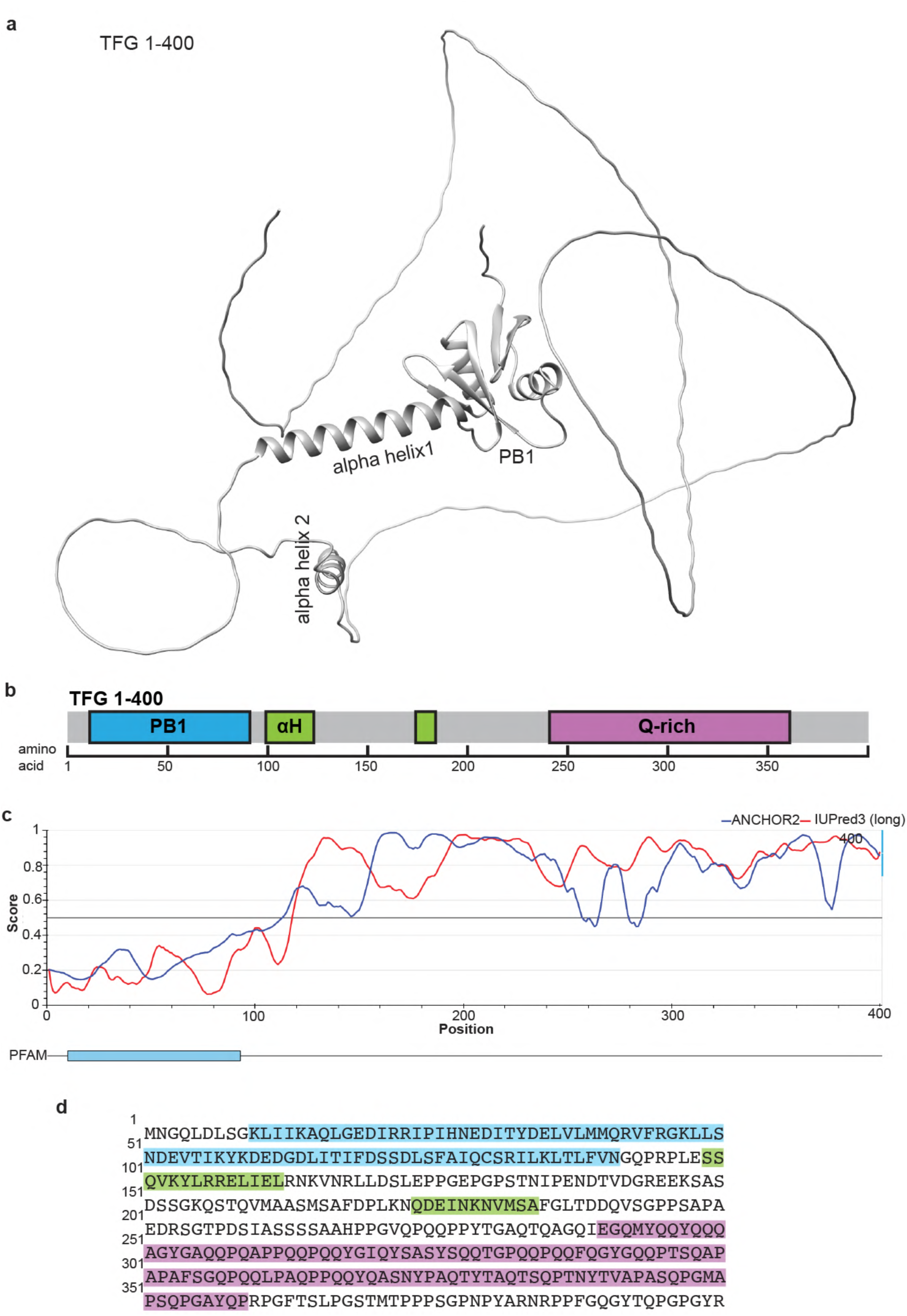
AlphaFold-predicted structure of TFG residues 1-400. IUPred predictions of full length TFG show regions of disorder. **a** Structure of TFG as predicted by AlphaFold with PB1, and high-confidence alpha helices 1 and 2 annotated. **b** Diagram of TFG domain structure (to scale). **c** IUPred3 predictions of TFG, with the red line symbolizing propensity for disorder in the structure and the blue line indicating the propensity to be part of a disordered binding region (58). **d** Amino acid sequence of TFG in rows of 50 amino acids, color-coded as in **b**.

**Extended Data Fig. 6.**
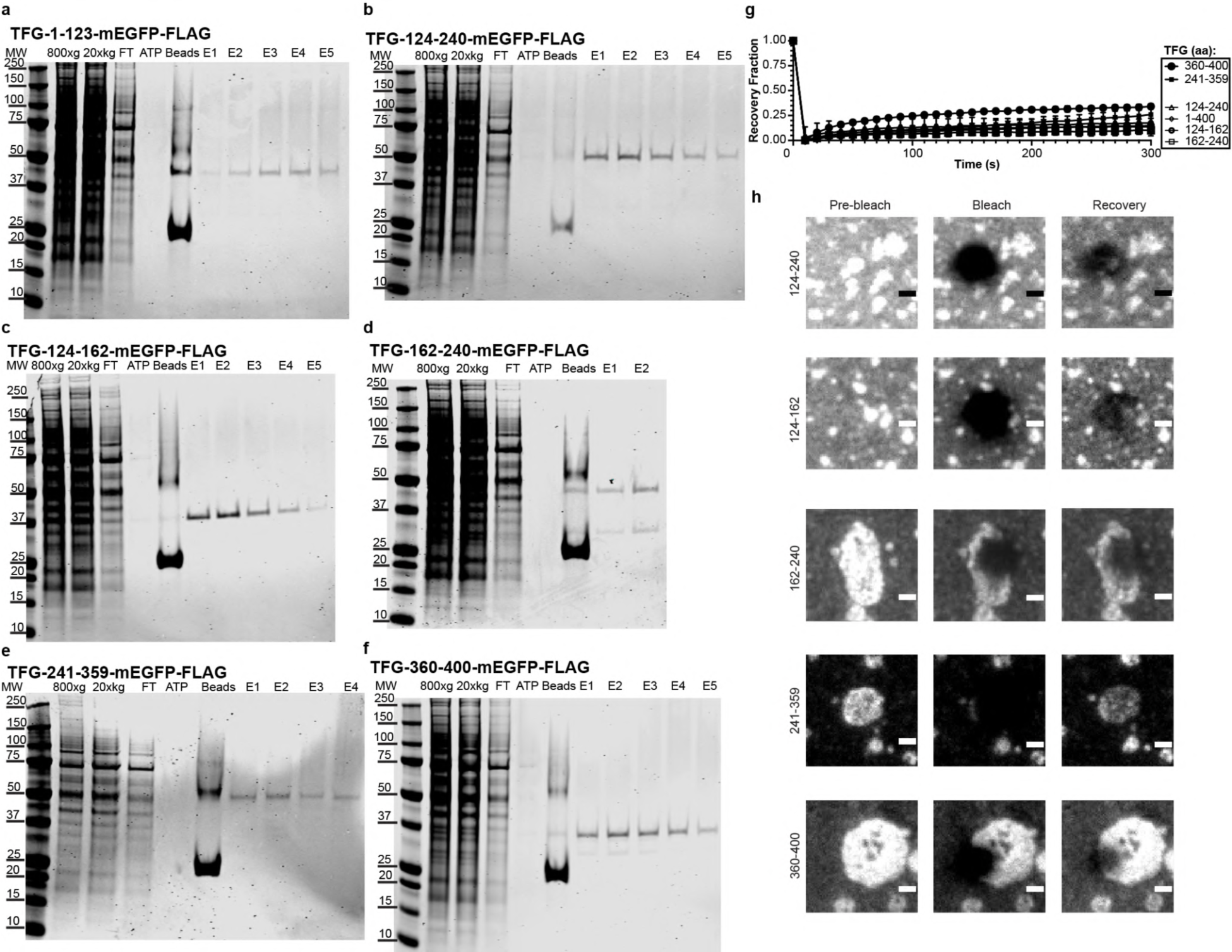
Purification overview and FRAP of truncated TFG constructs. Representative purification overview of **a** TFG-1-123-mEGFP-FLAG. **b** TFG-124-240-mEGFP-FLAG. **c** TFG-124-161-mEGFP-FLAG. **d** TFG-162-240-mEGFP-FLAG. **e** TFG-241-359-mEGFP-FLAG. **f** TFG-360-400-mEGFP-FLAG. **g** FRAP curves of condensates formed by TFG truncations. TFG 1-400: n = 8; TFG 124-240: n = 11; TFG 124-161: n = 10; TFG 162-240: n = 10; TFG 241-359: n = 15; TFG 360-400: n = 4. **h** Visual correlates for FRAP experiments of: TFG-124-240-mEGFP-FLAG, TFG-124-162-mEGFP-FLAG, TFG-162-240-meGFP-FLAG, TFG-241-359-mEGFP-FLAG, TFG-360-400-mEGFP-FLAG (HEPES/KOH pH 7.3; 150 mM KCl; 20% (v/v) PEG 8 kDa). Scale bars 1 μm. Larger, sponge-like condensates were selected to obtain partial FRAP data due to the size of individual condensates.

**Extended Data Fig. 7.**
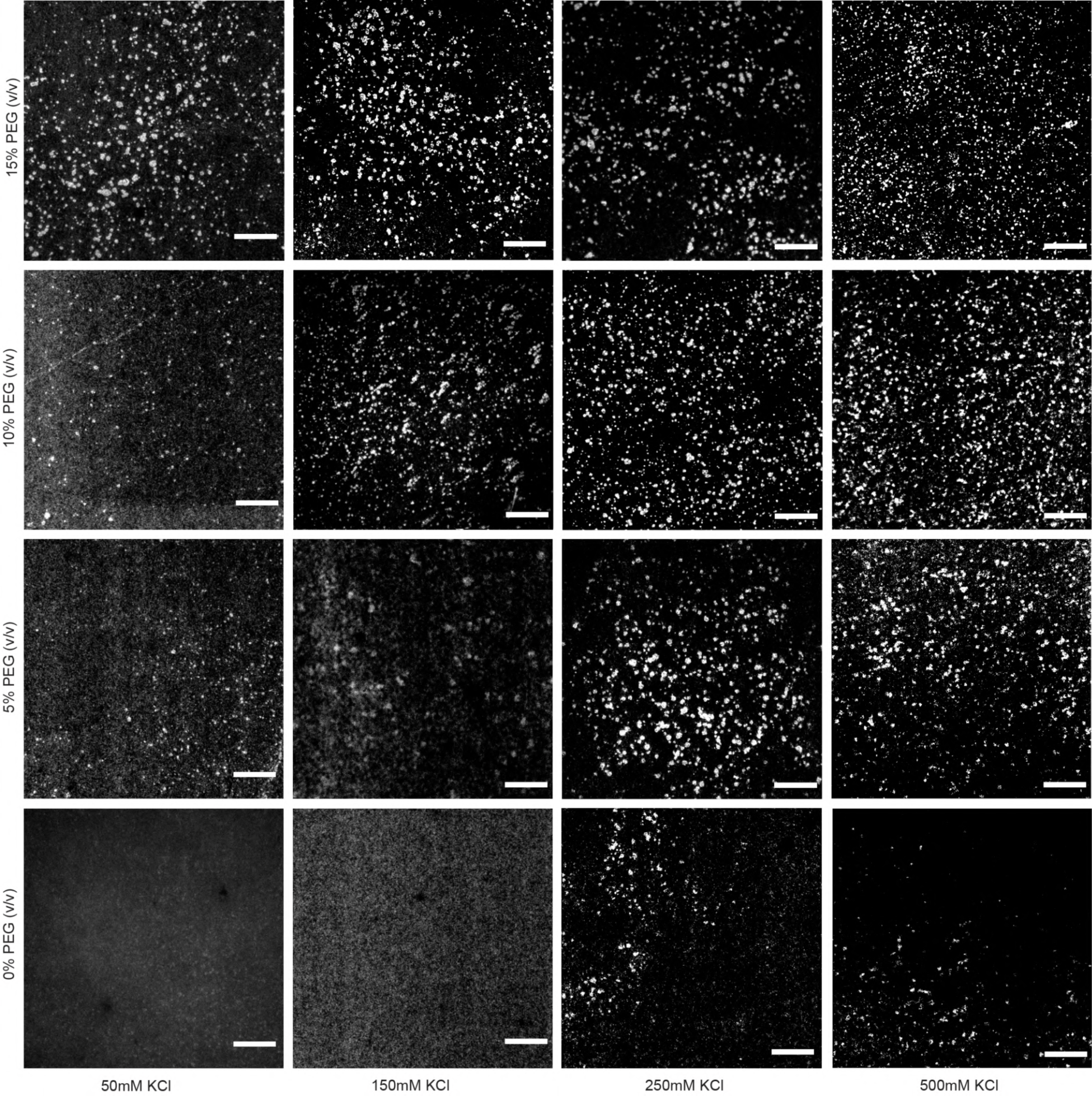
TFG phase diagram (visual correlates). Elevated buffer ionic strength promotes anisotropic condensation of TFG. TFG-mEGFP-FLAG: 70 nM; HEPES/KOH 7.3, X mM as indicated; X% (v/v) PEG 8 kDa as indicated. Scale bar 10 µm.

**Extended Data Fig. 8.**
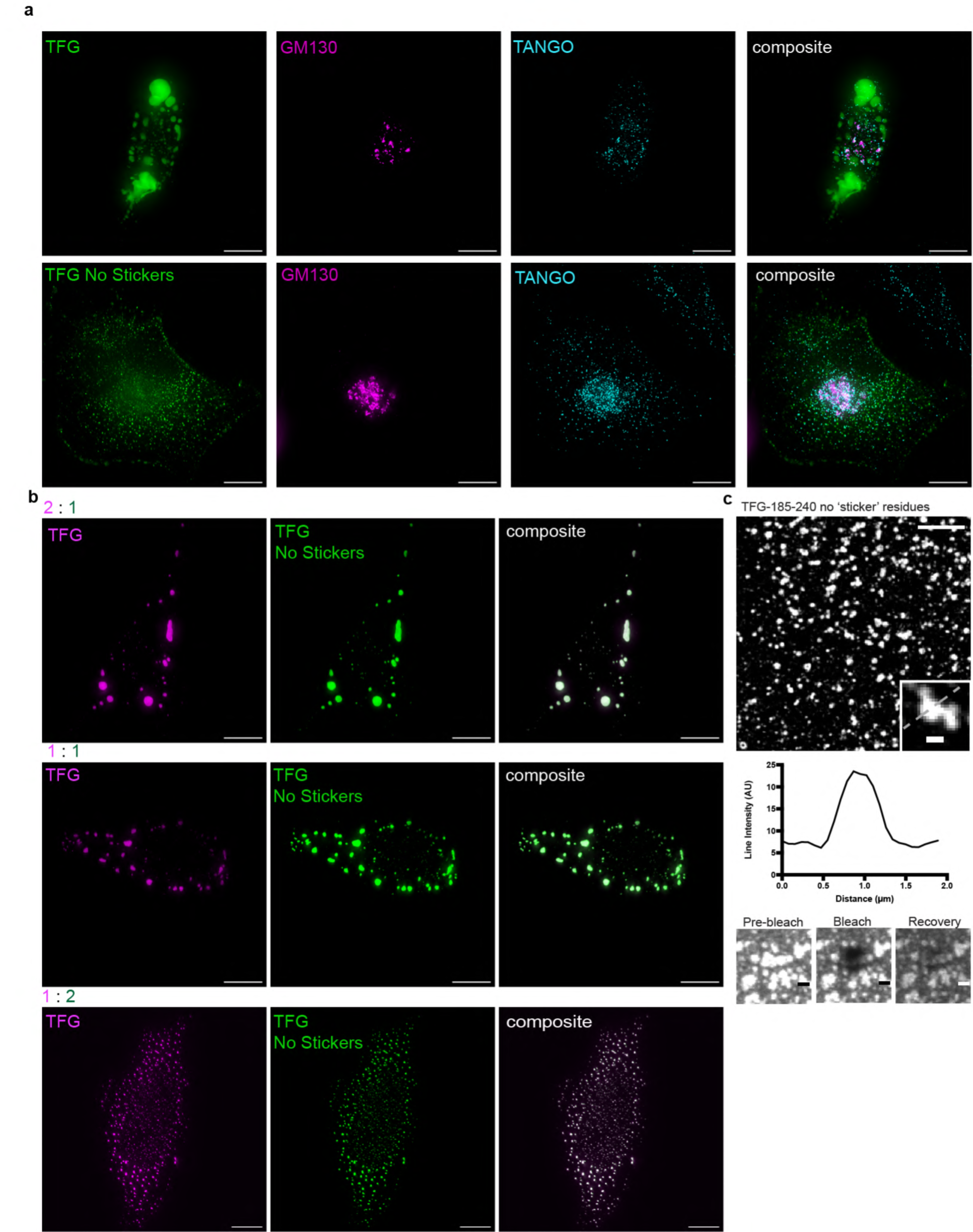
TFG disrupts endogenous ERES and Golgi markers only if ‘sticker’ residues are present. **a** Max. intensity Z-projections of HeLa cells transfected with TFG-mEGFP-FLAG or TFG-w/o ‘stickers’-mEGFP-FLAG and immunolabeled with GM130 and TANGO1 exhibiting Golgi disruption without sticker residues. Scale bar 10 µm. **b** Max. intensity Z-projections of HeLa cells transfected with FLAG-TFG-SNAP (stained with SiR) and TFG-w/o ‘stickers’-mEGFP-FLAG in ratios (as µg of plasmid DNA transfected per 100,000 cells) as indicated. Scale bar 10 µm. **c** Recombinant TFG-185-240-w/o ‘stickers’-mEGFP-FLAG (HEPES/KOH pH 7.3; 150 mM KCl; 20% (v/v) PEG 8 kDa). Scale bar 5 μm, 500 nm insets. Line intensity graph of condensate, represented by dashed line, shows no lumen. FRAP of TFG-185-240-w/o ‘stickers’-mEGFP-FLAG. Recovery t = 10 minutes. Scale bars 1 μm.

**Extended Data Fig. 9.**
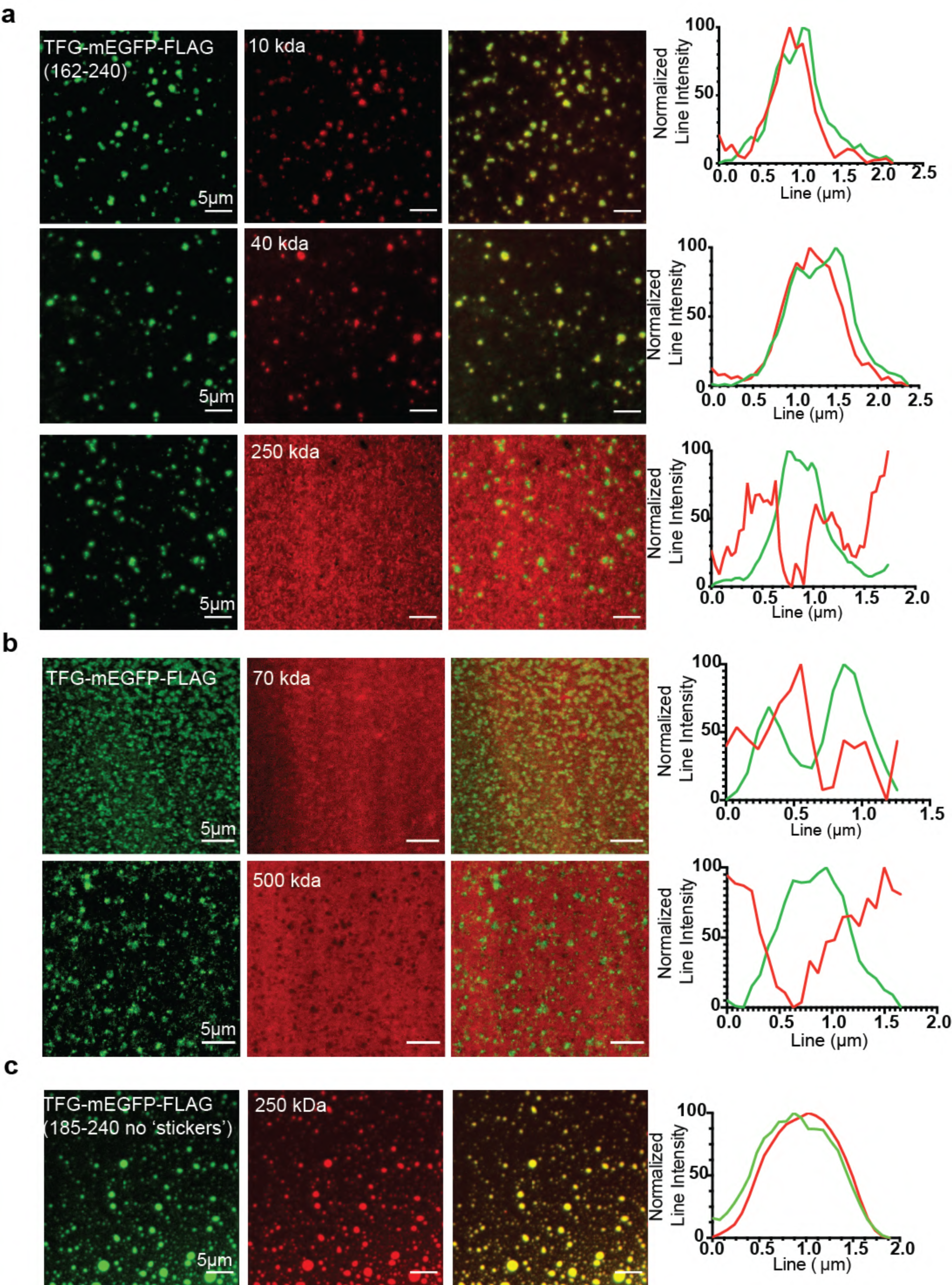
Visual correlates for TFG condensates exposed to dextrans of defined molecular weights. Recombinant protein condensates incubated for 10 minutes with dextran-TMR species as indicated. Normalized line intensities for representative images are given. Scale bars 5 µm. **a** TFG-162-240-mEGFP-FLAG. Dextran-TMR species of 10 kDa, 40 kDa, 250 kDa. **b** TFG-mEGFP-FLAG. Dextran-TMR species of 70 kDa and 500 kDa. **c** TFG-185-240-w/o ‘stickers’-mEGFP-FLAG. Dextran-TMR species of 250 kDa.

**Extended Data Fig. 10.**
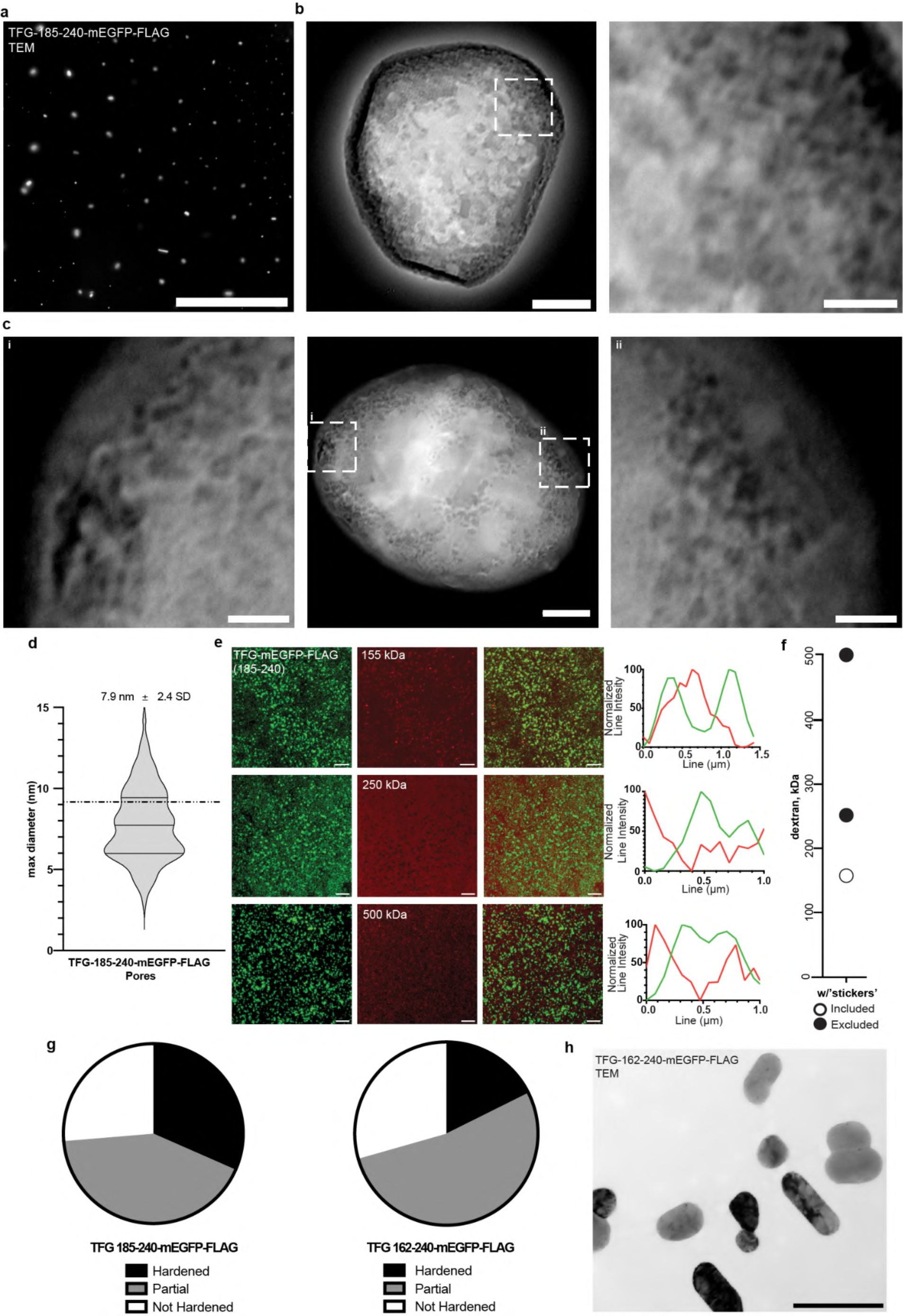
The alphahelical region (174-184) stabilizes TFG condensates but does not impact the condensation mechanism or alter condensate selectivity. **a** TEM overview of TFG-185-240-mEGFP-FLAG without negative staining. Scale bar 10 μm. **b** TEM micrograph of a single condensate of TFG-185-240-mEGFP-FLAG. Scale bar 100 nm. Magnification scale bar 25 nm. **c** TEM micrograph of a TFG-185-240-mEGFP-FLAG condensate. Scale bar 100 nm. Magnification scale bars: 25 nm. **d** Pore diameter quantification measured over the widest distance of pore. Dashed line represents average pore diameter of TFG 162-240-mEGFP-FLAG. n = 114 **e** Recombinant TFG-185-240-mEGFP-FLAG condensates incubated for 10 minutes with dextran-TMR species of 155 kDa, 250 kDa, and 500 kDa as indicated. Normalized line intensities for representative merged images are given. Scale bar 5 µm. **f** Plot representing dextran species included/in lumen of TFG-185-240-mEGFP-FLAG condensates: white-filled circles; excluded from TFG-185-240-mEGFP-FLAG condensates; back circles; permeability cutoff: 250 kDa. **g** Pie charts showing ratio of hardened, partially hardened, and not hardened condensates in TEM of TFG-185-240-mEGFP-FLAG (n = 19) and TFG-162-240-mEGFP-FLAG (n = 17) as indicated. **H** TEM overview of TFG-185-240-mEGFP-FLAG without negative staining. Scale bar 500 nm.

## Materials and Methods

### Cell Culture/Transfection/Labeling

HeLa cells (ATCC, CCL2) were grown at 37°C at 5% CO_2_ in Dulbecco’s modified eagle medium (Gibco 10566-016) supplemented with 10% FBS (Gibco A31604-01). Cells were seeded on glass-bottom Mattek dishes (P35GC-1.5-14-C) for live cell imaging or fixed with 4% PFA (Electron Microscopy Sciences, cat # 50980487) for 15 minutes. Fixed cells were then washed with PBS and permeabilized with permeabilization buffer (0.3% IGEPAL, 0.05% Triton-X 100, 0.1% BSA), then washed with wash buffer (0.05% IGEPAL, 0.05% Triton-X 100, 0.2% BSA), and blocked with blocking buffer (0.05% IGEPAL, 0.05% Triton-X 100, 5.0% normal goat serum) for 1 hour at room temperature. As indicated, cells were labeled with anti-GM130 (BD Biosciences, cat # BDB610822), anti-SEC16 (Invitrogen, cat # PA552182), anti-TFG (Abcam, cat # 156866), anti-TANGO1 (Abcam, cat # 244506) for 1 hour, washed, and labeled with goat-anti-mouse (Invitrogen, cat # A32727) and goat-anti-rabbit (Invitrogen, cat # A32732) Alexa Fluor Plus - conjugated secondary antibodies for 1 hour, subsequently washed and imaged. For ER-Golgi interface diameter measurements (Fig. 1g), n > 3 cells were utilized with multiple measurements derived from each.

For overexpression of TFG (Fig. 2, 3), cells were transfected with TFG-mEGFP-FLAG at 1 µg/µL using Fugene (Promega, cat # PRE2311) and left to express for 24 hours. 4-color micrographs (Fig. 1a) were obtained from HeLa cells co-transfected with mGFP-Sec16L at 3 µg/µL and FLAG-TFG-SNAP at 1 µg/µL and subsequently labeled with SNAP-Cell 647-SiR (NEB, S9102S) and immunostained with anti-GM130, anti-TANGO1, goat-anti-mouse Alexa Fluor Plus 405 (Invitrogen, cat # A48255), and goat-anti-rabbit Alexa Fluor Plus 555. Additional antibodies used in this study include: anti-GFP (Sigma, A11122) and anti-FLAG (Sigma, F1804).

For growth curve experiments, an siRNA transfection complex was prepared using TFG siRNA (L-016366-00-0005, Horizon Discovery Biosciences Limited). Oligonucleotides were diluted to 20 µM in ddH_2_O, and 10 µL were mixed into 175 µL of Opti-MEM (for mock knockdown, siRNA was omitted). Next, 3 µL of oligofectamine (Cat# 12252-011) was mixed into 12 µL of Opti-MEM. Both solutions were incubated for 5-10 minutes. Solutions were combined, incubated for 20 minutes, and 800 µL of Opti-MEM was added for a total of 1 ml of transfection mixture. 500,000 cells were spun at 200 x g, resuspended in 1 ml of transfection mixture, added to target cells for 20 hours in a 6 well plate. After incubation, cells were spun down at 200 x g, and resuspended in 2 ml of Expi expression medium. An automatic cell counter (Countess II; Thermofisher) was employed to analyze cell counts at indicated time points.

### Endogenous Tagging of TFG in HeLa Cells

TFG::mClover-FLAG knock-in HeLa cells were generated by ExpressCells (ExpressCells, Inc., Philadelphia, PA, USA) via CRISPR/Cas9, and validated by PCR, Sanger sequencing, and by fluorescence microscopy.

### Microscopy

Conventional (widefield/deconvolution, confocal) and super-resolution microscopy was performed on a Cytiva OMX SR microscope setup equipped with an Olympus PlanApo N 60X/1.42 oil objective. Multichannel alignment was performed by the manufacturer using TetraSpeck beads. Multicolor micrographs were aligned employing OMX-specific software packages (softWoRx). Images of immunolabeled cells (Extended Data Fig. 1a, b, e) were obtained using widefield microscopy with subsequent deconvolution via OMX-specific software packages (softWoRx). For time-lapse imaging of transfected HeLa cells (Extended Data Fig. 3a), images were acquired once every minute for 60 minutes using conventional widefield mode. Images of sponge-like TFG condensates (Extended Data Fig. 4b) were obtained via 3D structured illumination microscopy (3D-SIM), and images were reconstructed employing OMX-specific software packages (softWoRx).

SEM images (Fig. 4a) were taken on an FEI Quanta 450 with working distance of 10 mm, 5-10 kV, aperture 7. Sample was blotted on 0.1 µm filters, mounted on 12 mm Al stubs (Ted Pella, Redding, CA, cat# 16111) using double sticky carbon tabs. Samples were coated with 6 nm platinum using a Quora 150 V coater. TEM images (Fig. 4a, Extended Data Fig. 10a, b, c, h) were taken on an FEI Tecnai T12, 120 kV, with AMT digital camera. Sample was blotted onto 100 mesh carbon-formvar coated grids (Electron Microscopy Sciences, Hatfield, PA cat # FCF100H-Cu). All *in vitro* confocal, 3D-SIM, and TIRF images were taken with sample spotted on poly-D-lysine coated glass bottom dishes and at RT (Mattek, P35GC-1.5-14-C).

Diameter measurements (Fig. 1g, 2b, 3c) were taken using line scan measurements between pixels with highest brightness values. Heterozygous HeLa TFG-mClover-FLAG cells were seeded on glass-bottom Mattek dishes (P35GC-1.5-14-C) for live cell imaging with live imaging solution (Invitrogen A1429IDJ) supplemented with 10 mM final concentration of glucose (Sigma G8644-100ML). Cells were labeled with Hoechst 33342 (Invitrogen, cat # H3570) at a concentration of 5 μg/ml for 5 min. Micrographs of labeled cells (Fig 2a) were obtained using widefield microscopy.

Stimulated emission depletion (STED) images produced from HeLa cells seeded on glass-bottom Mattek dishes (P35GC-1.5-14-C) and transfected with TFG-mEGFP-FLAG at 1 µg/µL using Fugene (Promega, cat # PRE2311) for 24 hours. Cells were fixed with 4% PFA (Electron Microscopy Sciences, cat # 50980487) for 15 minutes and mounted with Prolong Diamond antifade mountant (Invitrogen cat # P36970). STED micrographs were obtained using an Inverted Leica DMi8 (UCSD Microscopy core) with 100x oil objective (excitation laser: 484 nm; depletion laser: 592 nm).

For FRAP analysis of TFG condensates in cells (Fig. 2), cells were transfected as described above and individual TFG condensates from n > 3 cells were either fully or partially bleached using the FRAP module in the OMX SR platform combined with confocal mode at 37°C. FRAP of *in vitro* TFG condensates from independent purifications was recorded at RT. Recovery was measured using the Time Series Analyzer V3 plugin in ImageJ^39^, and plots were generated using GraphPad Prism.

### Protein Purification from Expi293 Cells

Full-length and truncated TFG-encoding plasmids were transfected into Expi293F cells (ThermoFisher) at 1 µg/ml culture using Expifectamine 293 Reagent per manufacturer’s instructions. Time course experiments were conducted to ensure optimal expression time for constructs, with expression times varying from 16 hours to 72 hours. Expi293F cells were incubated at 37°C, 8% CO_2_ on a shaker at 120 rpm. Cells were pelleted at 800 x g for 10 minutes and frozen in liquid nitrogen. Pellets between 15 ml and 45 ml of culture, depending on the construct, were resuspended in 10 ml of buffer 1 [50 mM HEPES/KOH, pH 7.3, 150 mM KCl, EDTA-free protease inhibitor cocktail tablet (Roche)]. Cells were then homogenized employing 20 passages through a 25G needle. Lysate was centrifuged at 800 x g for 30 minutes. Pelleted material was discarded, and 5 ml buffer 2 [50 mM HEPES/KOH, pH 7.3, 1.2 M KCl, EDTA-free protease inhibitor tablet (Roche)] was added to supernatant to adjust salt concentration to 500 mM KCl final concentration. Salt adjusted lysate was then centrifuged at 20,000 x g for 15 minutes.

Pelleted material was discarded, and the supernatant was transferred to a conical tube containing 2.5 ml blocked FLAG-affinity resin equilibrated with buffer 3 [50 mM HEPES/KOH, pH 7.3, 500 mM KCl, EDTA-free protease inhibitor tablet (Roche)] and rotated for 1 hour at room temperature. The suspension was settled on a column that was equilibrated with 50 ml buffer 3. The resin was then washed with 100 ml buffer 3. Next, the column was drained and 1 ml buffer 4 [buffer 3 plus 0.5 mM ATP, 0.5 mM MgCl_2_ (Thermofisher)] was added to the resin and incubated for 10 minutes, followed by a wash of 50 ml buffer 3. If RNAse wash is indicated, 10 U of RNAse A (Thermofisher, Cat# 19101) was diluted into 1 ml total of buffer 3, added to the resin, incubated for 10 minutes, and followed by a wash of 50 ml of buffer 3. The proteins were eluted in buffer 5 [buffer 3 plus 200 μM FLAG peptide (Sigma-Aldrich)] for 1.25 hours, obtaining 5 fractions total. Elutions were analyzed on 4-20% Bis-Tris gradient gels, stained with Coomassie, and analyzed on a LI-COR Odyssey infrared scanner. After elution, fractions were frozen in liquid nitrogen before further processing. Quantitative Western blot analysis was employed to determine concentration of recombinant protein using a recombinant GFP standard (Roche, 11814524001) with a defined concentration of 1 mg/ml. Comparative Coomassie staining in conjunction with background-subtracted quantification of band intensity (LI-COR) of standards was employed to fit a gel-specific standard curve used to determine the concentration of a dilution series of candidate proteins analyzed on the same gel.

Concentration of samples was performed using a 50 kDa Amicon Ultra-0.5 Centrifugal Filter Unit (Millipore, UFC5050). The elution was thawed at 37°C with occasional vortexing, and the sample was centrifuged at 21.1 x g for 10 minutes before concentration. The column was washed in 200 μl steps with 2 ml total of 50 mM HEPES/KOH, pH 7.3 and 150 mM KCl, yielding a final 150 mM KCl final in the sample. Next, the sample was concentrated further stepwise in the desalting column, crowding agents and reductive agents were added as indicated, and the sample was used immediately for experiments without further storage.

### Probing Size Selectivity of TFG Condensates with Dextrans

Purified TFG-mEGFP-FLAG (+truncations) in 50 mM HEPES/KOH, pH 7.3 and 150 mM KCl was pipetted onto standard glass-bottom MatTek dishes (P35GC-1.5-14-C) to form condensates (with 20% v/v PEG 8 kDa as needed) and overlaid with fluorescent dextrans of various sizes at 10 kDa (Invitrogen D1816), 40 kDa (Invitrogen D1842), 70 kDa (Invitrogen D1819), 155 kDa (T1287-100MG, SIGMA-ALDRICH, INC.), 250 kDa (TMR-dex 250 kDa, Fina Biosolutions), and 500 kDa (52194-1G, SIGMA-ALDRICH, INC.) suspended in 50 mM HEPES/KOH, pH 7.3 and 150 mM KCl. Dextran species were added at a final concentration of 1 µM and incubated for 10 minutes. All images were taken in confocal mode using the OMX Flex SR microscope.

### RUSH Assay

HeLa cells (ATCC, CCL2) were seeded onto glass-bottom MatTek dishes (P35GC-1.5-14-C) for 24 hours and subsequently transfected with TFG siRNA (L-016366-00-0005, Horizon Discovery Biosciences Limited) and mock siRNA (D-001810-01-05, Horizon Discovery Biosciences Limited). 48 hours post knockdown, dishes were transfected with 1 µg of Str-KDEL_SBP-EGFP-ECadherin plasmid and left to express for an additional 24 hours. Cells were then imaged on the OMX SR platform in confocal mode at 37°C, and in HEPES-buffered live-cell imaging solution (Invitrogen A14291DJ) supplemented with 5 mM glucose. To release RUSH cargo from the ER, biotin (B4501, SIGMA ALDRICH, INC.) was added to a final concentration of 100 µM, and cells were imaged at 5-minute intervals for 60 minutes. Statistical significance was determined via two-tailed, unpaired t-tests: ***: p < 0.001, **: p<0.01.

### Plasmids

The sequences for plasmids used were sourced from UniProtKB/Swiss-Prot and included: TFG Q92734. The fragments for constructs used were generated via commercial gene synthesis (gBLOCK; IDT) and contained the sequence for mEGFP and FLAG. gBLOCKs and the pSNAP_f_ plasmid (NEB, N9183S) were both digested with NheI (NEB, R0189S), removing SNAP and inserting the commercially obtained gene via ligation with T4 ligase (Sigma, 10481220001) per manufacturer’s instructions. TFG-185-240-w/o ‘stickers’-mEGFP-FLAG refers to TFG-185-240-(ΔF185, ΔL187, Δ192, ΔI210, ΔV222, ΔI239)-mEGFP-FLAG. TFG-360-400-mEGFP-FLAG was generated via PCR using primers FWD:5’-tatatagctagcATGAGACCAGGTTTTACTTCACTTCCT-3’ and REV:5’-GATTACAAGGATGACGACGATAAGctcgaggttaat-3’. The amplification product was gel-purified using Qiaquick Gel Extraction Kit (Qiagen, 28706X4), digested with NheI and XhoI (NEB, R0146S), and inserted into a similarly digested pSNAP_f_ plasmid using T4 ligase. The ligation mixture was transformed into Subcloning Efficiency DH5alpha competent cells (ThermoFisher, 18265017), plated onto 100 μg/ml ampicillin agar plates (Biomyx, LA-2100), and several single colonies were cultured in LB and 100 μg/ml ampicillin. Single colony derived cultures were subsequently mini-prepped using a Qiaprep Spin Miniprep Kit (Qiagen, 27104), and plasmid sequences were confirmed via sequencing by Azenta/GeneWiz. pmGFP-Sec16L was a gift from Benjamin Glick (Addgene plasmid # 15776; http://n2t.net/addgene:15776; RRID:Addgene_15776). Str-KDEL_SBP-EGFP-Ecadherin was a gift from Franck Perez (Addgene plasmid # 65286; http://n2t.net/addgene:65286; RRID:Addgene_65286).

### Condensate Thermoresponsiveness Assay in Live Cells

100,000 CCL2 HeLa cells were seeded onto 35 mm glass-bottom Mattek dishes and allowed to adhere to plate overnight. After 24 hours, cells were transfected with 1 μg TFG-mEGFP-FLAG using the transfection reagents (Fugene + Optimem) per manufacturer’s instructions and subsequently incubated in DMEM + 10 % FBS. 24 hours post-expression of TFG-mEGFP-FLAG, Mattek dishes were removed from the incubator the medium was exchanged for 1ml live-cell imaging buffer pre-incubated to defined temperatures, incubated for 5 min, and subjected to microscopy on an OMX DeltaVision SR setup in confocal mode. The axial dimensions of condensates for a given temperature were quantified to calculate average condensate circularity (ratio of width divided by length) per temperature condition.

